# SCD1 and monounsaturated lipids are required for autophagy and survival of adipocytes

**DOI:** 10.1101/2023.10.27.564376

**Authors:** Hiroyuki Mori, Sydney K. Peterson, Rachel Simmermon, Katherine A. Overmyer, Akira Nishii, Emma Paulsson, Ziru Li, Annie Jen, Romina M. Uranga, Jessica Maung, Warren T. Yacawych, Kenneth T. Lewis, Rebecca L. Schill, Taryn Hetrick, Ryo Seino, Ken Inoki, Joshua J. Coon, Ormond A. MacDougald

## Abstract

Exposure of adipocytes to ‘cool’ temperatures often found in the periphery of the body induces expression of Stearoyl-CoA Desaturase-1 (SCD1), an enzyme that converts saturated fatty acids to monounsaturated fatty acids. In this study, we employed *Scd1* knockout cells and mouse models, along with pharmacological SCD1 inhibition, to investigate further the roles of SCD1 in adipocytes. Our study reveals that production of monounsaturated lipids by SCD1 is necessary for fusion of autophagosomes to lysosomes and that with a SCD1-deficiency, autophagosomes accumulate. In addition, SCD1-deficiency impairs lysosomal and autolysosomal acidification resulting in vacuole accumulation and eventual cell death. Blocking autophagosome formation or supplementation with monounsaturated fatty acids maintains vitality of SCD1-deficient adipocytes. Taken together, our results demonstrate that *in vitro* inhibition of SCD1 in adipocytes leads to autophagy-dependent cell death, and *in vivo* depletion leads to loss of bone marrow adipocytes.

## INTRODUCTION

Temperatures at which cells function varies widely depending on location within the body. The core is typically maintained at around 37°C whereas the shell - composed of peripheral extremities and subcutaneous regions of the trunk - displays a broader and generally cooler temperature range. Notably, adipocytes in subcutaneous, bone marrow, and dermal depots primarily exist at temperatures below 37°C, contrasting with their visceral counterparts in the body’s core. In a previous study, we explored the adaptability of white adipocytes to temperature changes by subjecting them to 31°C - the ‘cool’ temperature under which distal bone marrow and subcutaneous adipose tissues often exist when animals and humans are in a 22°C environment ^1,2^. This cool exposure induced expression of Stearoyl-CoA Desaturase-1 (SCD1), an enzyme that catalyzes conversion of saturated fatty acids to monounsaturated fatty acids ^3^.

Although SCD1 is expressed at low levels in most cell types, expression is much higher in lipogenic tissues including liver and white adipose tissue (WAT) ^4^. Global SCD1 deletion in mice led to increased energy expenditure and insulin sensitivity ^5^. Reduced SCD1 activity in liver and adipose tissues was not sufficient to elicit the phenotypes of global *Scd1*KO mice (GKO) ^6–8^. Interestingly, skin-specific *Scd1* knockout mice had increased water and heat loss across the surface of the skin and thus recapitulated the hypermetabolic phenotype observed in global *Scd1*KO mice ^9^. We previously reported that SCD1 expression is regulated by temperature, with considerably more SCD1 protein observed in bone marrow and subcutaneous adipose tissues of rodents housed at 22°C compared to 29°C ^3^. Consistent with elevated SCD1, decreased amounts of saturated lipids and increased proportions of monounsaturated lipids were observed within triacylglycerols of bone marrow adipose tissue (BMAT) and gluteal WAT of rodents housed at 22°C ^3^.

SCD1 and monounsaturated fatty acids (MUFAs) are associated with several functions in adipose and other tissues. In addition to lipid and carbohydrate metabolism, SCD1 also plays roles in production of adipokines, insulin sensitivity, endoplasmic reticulum stress, cancer cell survival, and autophagy ^10–12^. Autophagy consists of forming double-membrane vesicles encapsulating protein aggregates, damaged organelles such as mitochondria, and bulk cytoplasm, which then fuse with lysosomes to degrade and recycle their contents ^13–15^. Although it is traditionally viewed as an adaptive process enabling cells to survive stresses like nutrient deprivation, increasing evidence suggests autophagy may mediate cell death during development and pathogenesis ^13,14,16^. However, significant gaps remain in our understanding of autophagy-dependent cell death (ADCD) ^13^. The interplay between *de novo* lipogenesis and autophagy has emerged as a fascinating area of study in cellular biology. Fatty acids produced by fatty acid synthase (FASN) are essential for the autophagy process including autophagosome and lysosome maturation and fusion in adipocytes ^17^. The relationships between lipogenesis and autophagy are fundamental cell biological processes in that acetyl-CoA carboxylase1 (ACC1), the rate-limiting step of fatty acid biosynthesis, promotes autophagy and survival in chronologically aging yeast ^18^. In addition, deficiency of the *Scd1* orthologue in yeast demonstrated that MUFAs are required for formation of autophagosomes ^10^. However, to what extent *de novo* lipogenesis and SCD1 contribute to the control of autophagy in adipocytes remains elusive.

To further investigate roles of SCD1 in adipocytes, particularly in relation to cool temperature adaptation, we employed a genetic approach using SCD1 knockout cells and mice, along with pharmacological inhibition of SCD1. Our observations did not suggest that SCD1-deficiency leads to beige adipocyte formation, as previously suggested ^19^, but instead pointed to SCD1 inhibition causing adipocyte death. Our exploration of cell death processes in SCD1-deficient adipocytes revealed that both pharmacological and genetic inhibition of SCD1 in adipocytes causes ADCD. Impaired fusion of autophagosomes to lysosomes, coupled with reduced lysosomal acidification in *Scd1*KO adipocytes, resulted in vacuole accumulation and ultimately, cell death. Our argument is substantiated by evidence that blocking autophagosome formation protects *Scd1*KO adipocytes from cell death, a result analogous to the cell survival observed after supplementation of SCD1-deficient adipocytes with MUFAs.

## RESULTS

### Beige adipocyte markers were not induced in *Scd1*KO adipocytes

SCD1 resides in the endoplasmic reticulum where it converts Coenzyme A-derivatives of palmitic (C16:0) and stearic (C18:0) acids to palmitoleic (C16:1, n-7) and oleic (C18:1, n-9) acids, respectively. We recently reported that exposure of cultured adipocytes to cool temperatures (31°C) increases expression of SCD1 (**Figure 1A**) ^3^. To determine whether increased monounsaturated fatty acids (MUFAs) under cool stimulation was dependent on SCD1, we performed lipidomic analyses of *Scd1* KO adipocytes (**Figure 1B and Figure S1**). Cool adaptation of control adipocytes increased the proportion of several MUFA-containing lipid species, including triacylglycerol (TAG), phosphatidylcholine (PC), phosphatidylinositol (PI), and phosphatidylethanolamine (PE), and these lipids were reduced in *Scd1*KO adipocytes at both 37°C and 31°C (**Figure 1A, 1B, and Figure S1**). In contrast, lipid species containing mainly saturated lipids were often suppressed with cool adaption and were more abundant in *Scd1*KO adipocytes (**Figure 1B, and Figure S1**). These data suggest that induction of SCD1 with cool adaptation increases lipid monounsaturation, and that SCD1 is the predominant SCD isoform in cultured adipocytes.

**Figure 1.**
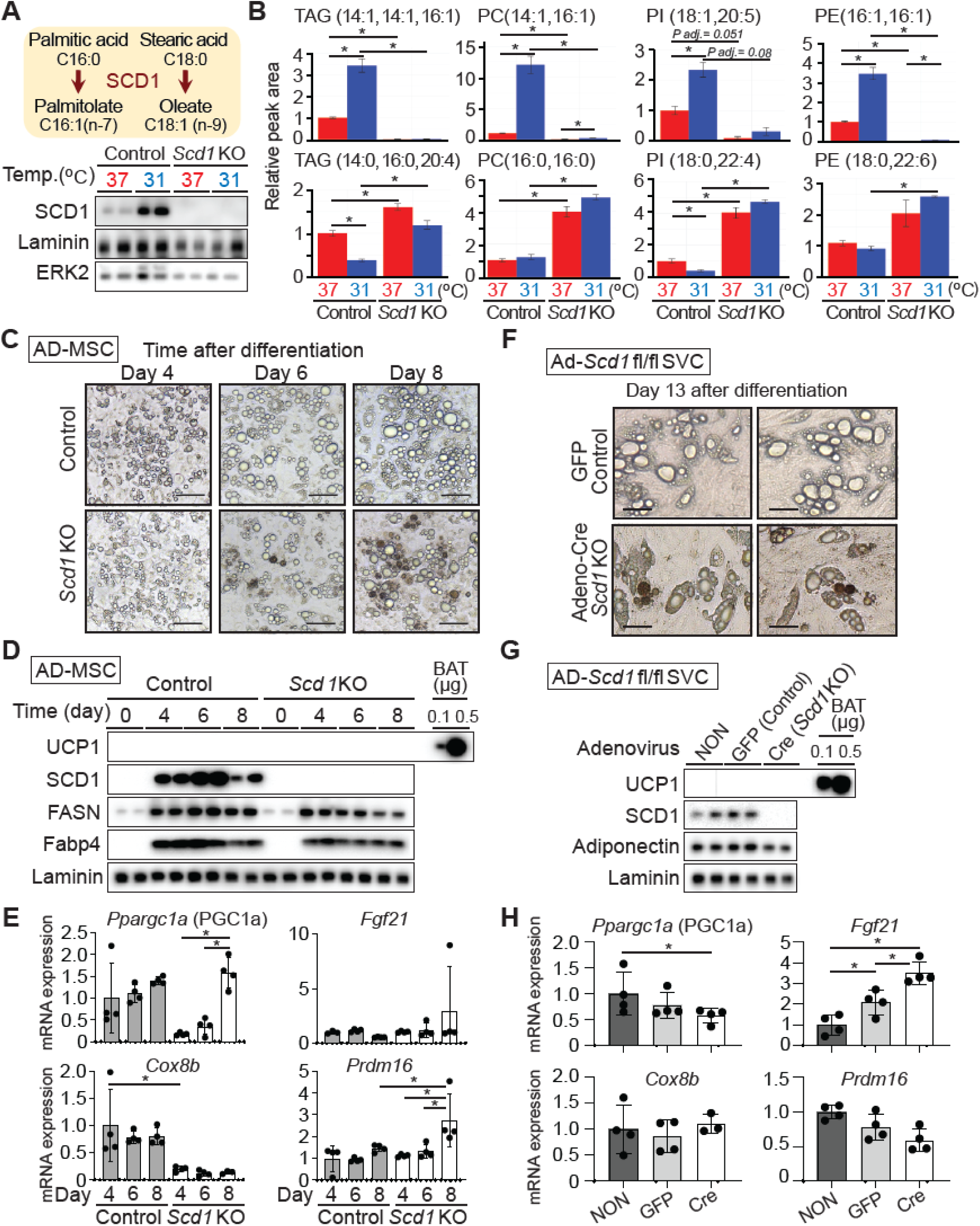
Beige adipocyte markers are not induced in *Scd1*KO adipocytes. (**A**) SCD1 converts saturated fatty acids (e.g., palmitic and stearic acids) to monounsaturated fatty acids (e.g., palmitoleic and oleic acids, respectively). SCD1 is induced at 31°C and is undetectable in *Scd1*-KO adipocytes. MSCs from control and global *Scd1* KO mice were cultured at 37°C prior to and for the first four days of adipogenesis, after which cells were moved to 31°C for 12 days. (**B**) Decreased monounsaturated lipids (upper panels) and increased saturated lipids (lower panels) in *Scd1*KO adipocytes. Adipocytes were cultured at 37°C or 31°C for 12 days. Peak area of each lipid is expressed as fold change relative to 37°C control (*n* = 3). Data are presented as mean ± SD. \**p* < 0.05. (**C**) Cultured *Scd1*KO adipocytes acquire brown morphological structures during terminal differentiation. Scale bar indicates 100 μm. (**D**) Undetectable UCP1 expression in *Scd1*KO adipocytes for the indicated days after differentiation. Eleven µg of MSC-derived adipocyte (AD-MSC) or 0.1 or 0.5 µg BAT lysate was evaluated by immunoblot for UCP1. Data is representative of 3 independent experiments. (E) Beige adipocyte markers were not induced in *Scd1*KO adipocytes except *Prdm16* at Day 8 (*n* = 4). Gene expression was normalized to geometric mean value of *Hprt*, *Gapdh*, *Ppia*, and was expressed relative to control at Day 4 (*n* = 4). \**p* < 0.05. Data is representative of 2 independent experiments. (**F**-**H**) Differentiated adipocytes from *Scd1*^fl/fl^ stromal vascular cells (SVCs) were treated with adenoviral GFP or adenoviral Cre recombinase at Day 4 of differentiation. (**F**) Morphology of *Scd1*KO adipocytes is characterized by aggregations of brown structures. Scale bar indicates 50 μm. (**G**) SCD1 and UCP1 protein expression in adipocytes following adenoviral GFP or Cre infection. Eleven µg of adipocyte or 0.1 or 0.5 µg BAT lysate was evaluated by immunoblot for UCP1. (**H**) Beige adipocyte markers were not induced in *Scd1*KO adipocytes except *Fgf21* (*n* = 4). Gene expression was normalized to geometric mean value of *Hprt*, *Gapdh*, *Ppia*, *Rpl32*, and was expressed relative to non-infected adipocytes (*n* = 4). \**p* < 0.05.

It was reported that cultured SCD1 knockout cells are prone to beiging and have impaired lipid accumulation during adipogenesis ^19^. Consistent with this observation, we observed that some *Scd1*KO mesenchymal stem cells (MSC) adipocytes and Cre-infected *Scd1*^fl/fl^ adipocytes acquire a brown morphology characterized by an aggregation of compact brown structures during terminal differentiation (**Figure 1C, 1F**, and **Figure S2A**). We performed immunoblot analyses to confirm that this morphology is related to beige adipogenesis and observed no UCP1 induction in *Scd1*KO adipocytes (**Figure 1D** and **1G**). Furthermore, beige adipocyte markers, including *Ppargc1a*, *Fgf21*, *Cox8b* and *Prdm16* were not consistently induced in *Scd1*KO adipocytes (**Figure 1E** and **1H**). Taken together, these data indicate that although SCD1 deficiency caused aggregations of brown structures reminiscent of beige adipocytes, these cells did not acquire molecular characteristics of beiging under our experimental conditions.

### Brown structures in *Scd1*KO MSC adipocytes are features of cell death

We next investigated whether brown structures newly arose or were transformed from existing cells. Time-lapse microscopy revealed that brown aggregates formed on lipid droplets of differentiating adipocytes (**Figure 2A**). This morphological change occurred rapidly, and brown structures formed within a 2-hour duration of imaging. To evaluate whether adipocytes containing brown structures were viable or dead, we first observed that they were efficiently labeled with Trypan blue (**Figure 2B**), suggesting that they were dead. To confirm cell death by another method, Calcein-AM and ethidium homodimer (EthD-1) dyes were used. Whereas almost all control adipocytes were stained with Calcein-AM as a live cell marker, fewer cells were Calcein-AM positive in *Scd1*KO adipocytes (**Figure 2C**). Increased frequency of EthD-1 positive nuclei, an indicator of dead and dying cells, was observed in *Scd1*KO adipocytes. EthD-1 positive nuclei were also frequently observed in *Scd1*KO adipocytes not stained with Calcein-AM and containing brown structures (**Figure 2C**).

**Figure 2.**
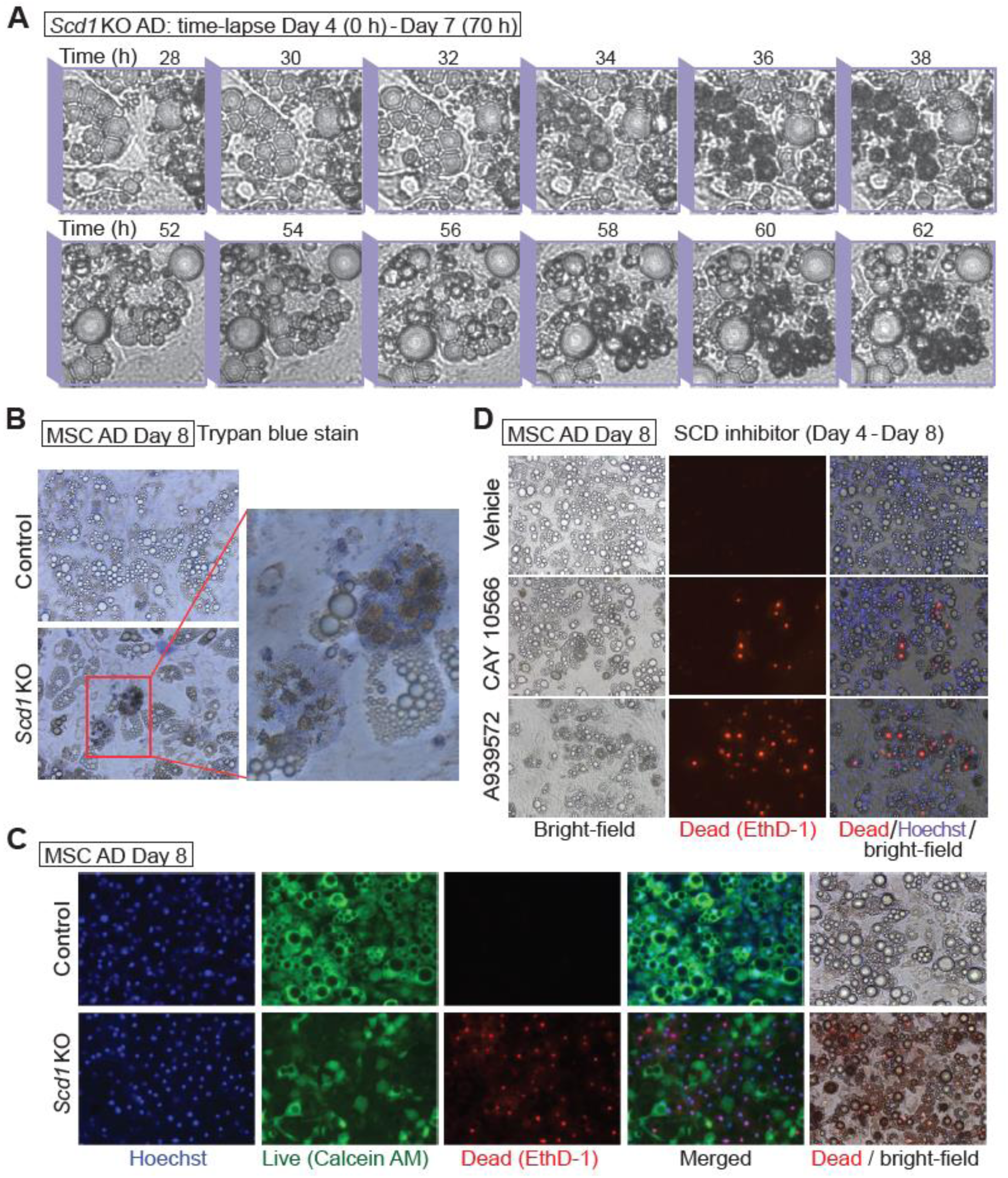
Brown structures in *Scd1*KO MSC adipocytes are features of cell death. (**A**) Representative pictures of time-lapse recording for *Scd1*KO MSC adipocytes acquiring brown structures. Cultured *Scd1*KO adipocytes were monitored every 2 h from Day 4 (0 h) to Day 7 (70 h). Difficulty with maintenance of focus during data acquisition resulted in some lipid droplets appearing to have double membranes. (**B**) *Scd1*KO adipocytes with brown structures are dead, and stain positive with trypan blue. Adipocytes were stained with Trypan blue at Day 8 after differentiation. (**C**) Staining with cell death indicator, EthD-1, was observed in *Scd1*KO adipocytes. Adipocytes cultured in DMEM/F12 with 2% FBS were stained with a live cell marker Calcein AM, dead cell marker EthD-1, and Hoechst at Day 8 after differentiation. (**D**) Adipocyte death caused by pharmacological inhibition of SCD1. Adipocytes were cultured with SCD inhibitors: 30 nM CAY10566 or 30 nM A-939572 for 4 days, then stained EthD-1, and Hoechst. Data shown in (**B**)-(**D**) are representative of at least 3 independent experiments.

Next, to examine whether pharmacological inhibition of SCD1 induced cell death in cultured adipocytes, MSCs were differentiated for 8 days, then treated with SCD inhibitors for 4 days. All three SCD1 inhibitors (CAY10566, A-939572, and MF-438) at a concentration of ∼30 nM induced brown structures (**Figure S2B**) that appeared similar to morphological changes observed adipocytes genetically deficient for SCD1. This phenomenon was also evident in adipocytes derived from human SVCs treated with an SCD1 inhibitor (**Figure S2C**). Consistent with the *Scd1*KO adipocytes, brown structure formation induced by SCD1 inhibitors also displayed EthD-1 positive nuclei (**Figure 2D**), suggesting pharmacological inhibition of SCD1 also induces cell death in cultured adipocytes. Taken together, these results indicate that pharmacological inhibition and genetic deficiency of SCD1 cause adipocyte death *in vitro*.

### MUFA supplementation rescues adipocytes from cell death induced by SCD1 deficiency

Pharmacological and genetic inhibition of SCD1 causing adipocyte death led us to investigate next whether MUFAs, the enzymatic products of SCD1, protect adipocytes against the formation of brown structures and cell death. In *Scd1*KO adipocytes, supplementation with 100 µM palmitoleic (C16:1) or oleic (C18:1) acids, or a mixture of both MUFAs (50 µM each) from day 4 to 8 of differentiation blocked the appearance of brown structures (**Figure 3A-3C**). Whilst addition of palmitic acid (C16:0) to *Scd1*KO adipocytes did not rescue these phenotypes, it caused formation of additional brown structures compared to BSA-treated cells (**Figure 3A-3C**). Brown-colored structures were less stained with BODIPY or Oil Red O than normal lipid droplets (**Figure 3B** and **Figure S2A**). There were no identified toxic signs attributed to this concentration of non-esterified fatty acids in control adipocytes. To determine whether MUFAs also rescue cell death induced by SCD1 inhibitors, adipocytes were cultured with SCD1 inhibitors with or without a 1:1 mixture of 50 µM palmitoleic and oleic acids. The number of brown structures induced by pharmacological inhibition of SCD1 was decreased with MUFA supplementation (**Figure 3D**). Taken together, these data suggest that conversion of saturated lipids to monounsaturated lipids is critical to protect adipocytes deficient for SCD1 from cell death.

**Figure 3.**
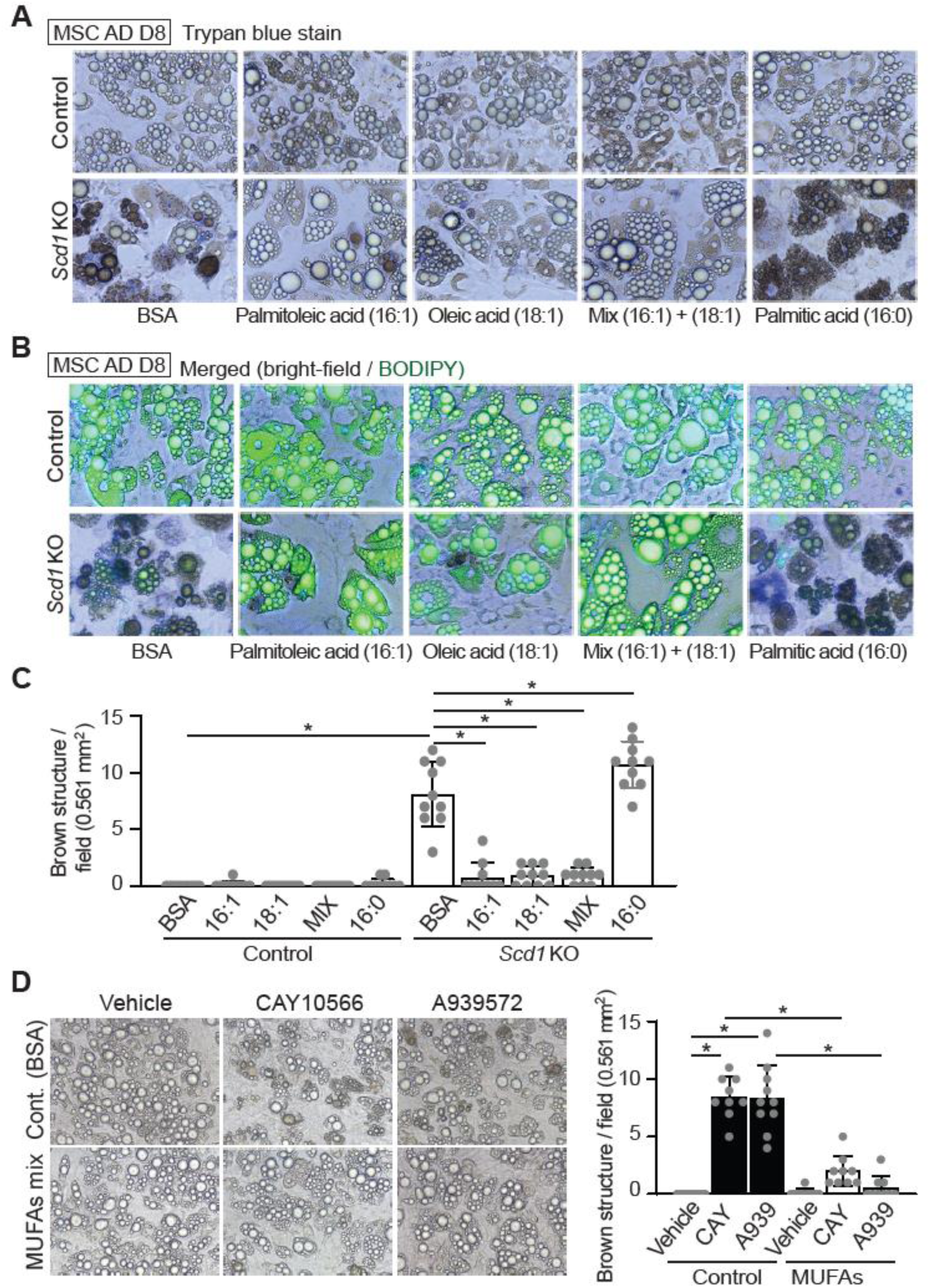
Supplementation of *Scd1*KO adipocytes with MUFAs maintains cell viability. (**A**-**C**) Supplementation with 100 µM palmitoleic (C16:1) or 100 µM oleic (C18:1) acids, or a mixture of 50 µM of each MUFA to *Scd1*KO adipocytes from day 4 to 8 of differentiation reduced appearance of brown structures. Addition of palmitic acid (C16:0) increased frequency of brown structures in *Scd1*KO adipocytes. (**A**) Phase contrast microscopy. (**B**) Fluorescent microscopy of cells following staining of neutral lipid droplets with BODIPY. (**C**) Quantification of brown structures observed per field. Brown structures were quantified from 10 fields in each group as shown in (**A**), with each field measuring 0.561 mm^2^. \**p* < 0.05. Data shown are representative of at least 3 independent experiments. (**D**) Supplementation with a MUFA mixture blocked adipocyte death following SCD1 inhibition. Adipocytes were cultured with SCD inhibitors - 30 nM CAY10566 or 30 nM A-939572 for 4 days with or without a mixture of MUFAs (50 µM of palmitoleic and oleic acids). Controls were treated with BSA alone. Brown structures were quantified from 9 fields in each group, with each field measuring 0.561 mm^2^. \**p* < 0.05.

### Decreased bone marrow adipose tissue (BMAT) volume and number in proximal and distal tibiae in bone marrow adipocyte (BMAd) specific SCD1 knockout mouse

To determine whether SCD1 is required for survival of adipocytes *in vivo*, we knocked *Scd1* out of BMAd, a cell type that expresses high levels of SCD1 when mice are housed at 22°C ^3^. BMAd-specific *Scd1* knockout (BMAd-*Scd1*KO) mice were generated by crossing *Scd1*^fl/fl^ mice with a BMAd-*Cre* mouse model ^20^. First, we validated deletion of SCD1 in caudal vertebrae, where constitutive bone marrow adipocytes (cBMAd) comprise ∼4% of total cellularity, and observed reduced expression of SCD1 in BMAd-*Scd1*KO mice (**Figure 4A**). No reduction in SCD1 was observed in WAT depots, where adipocytes are more plentiful (∼20%) than in caudal vertebra. Differentiated MSCs from control and *Scd1* KO mice were included as immunoblot controls (**Figure 4A**). To confirm the functional impact of SCD1 deletion in BMAT, we performed lipidomic analyses on distal tibia and caudal vertebra, both of which have significant amounts of BMAT. We focused our analyses on TAG and diacylglycerols (DAG) that are abundant in BMAds, rather than phospholipids, which are found in all cells and thus dilute specific effects of SCD1 deficiency in BMAds. We confirmed the expected enzymatic roles of SCD1 in BMAds, and generally observed lipids with decreased monounsaturated lipids and increased saturated lipids in BM-*Scd1*KO tibia and caudal vertebrae (CV) (**Figure S3**). No genotype-specific differences were observed in BMAd-*Scd1*KO mice, including body weight, length, lean or fat mass, weights of WATs, and glucose tolerance (**Figure S4A**-**S4F**). A slight increase in insulin sensitivity was observed in male BMAd-*Scd1*KO mice (**Figure S4G**), whereas there was no change in insulin sensitivity in female mice (**Figure S4H**). To quantify effects of SCD1-deficiency on bone marrow adiposity and cellularity, we used osmium tetroxide and μCT of tibia, and hematoxylin and eosin staining of BMAT sections. Regulated BMAT (rBMAT) volume was decreased in the proximal tibia of both sexes and constitutive BMAT (cBMAT) volume was reduced in distal tibia of male BMAd-*Scd1*KO mice (**Figure 4B-4D**). Histological analyses revealed decreased rBMAd number in proximal tibiae of BMAd-*Scd1*KO mice (**Figure 4E, Figure S5A**, and **S5B**). Depletion of SCD1 caused smaller size of cBMAd in distal tibia only in male mice (**Figure 4F-4H, Figure S5C-5E**). Although loss of BMAds has been shown to cause elevated bone mass ^21^, no differences were observed in bone volume fraction, bone mineral density, or other measure of bone mass in male and female BMAd-*Scd1*KO mice compared to controls (**Figure S5F-S5K**). The size of adipocytes in WAT was not affected in BMAd-*Scd1*KO mice (**Figures S5L-S5M**).

**Figure 4.**
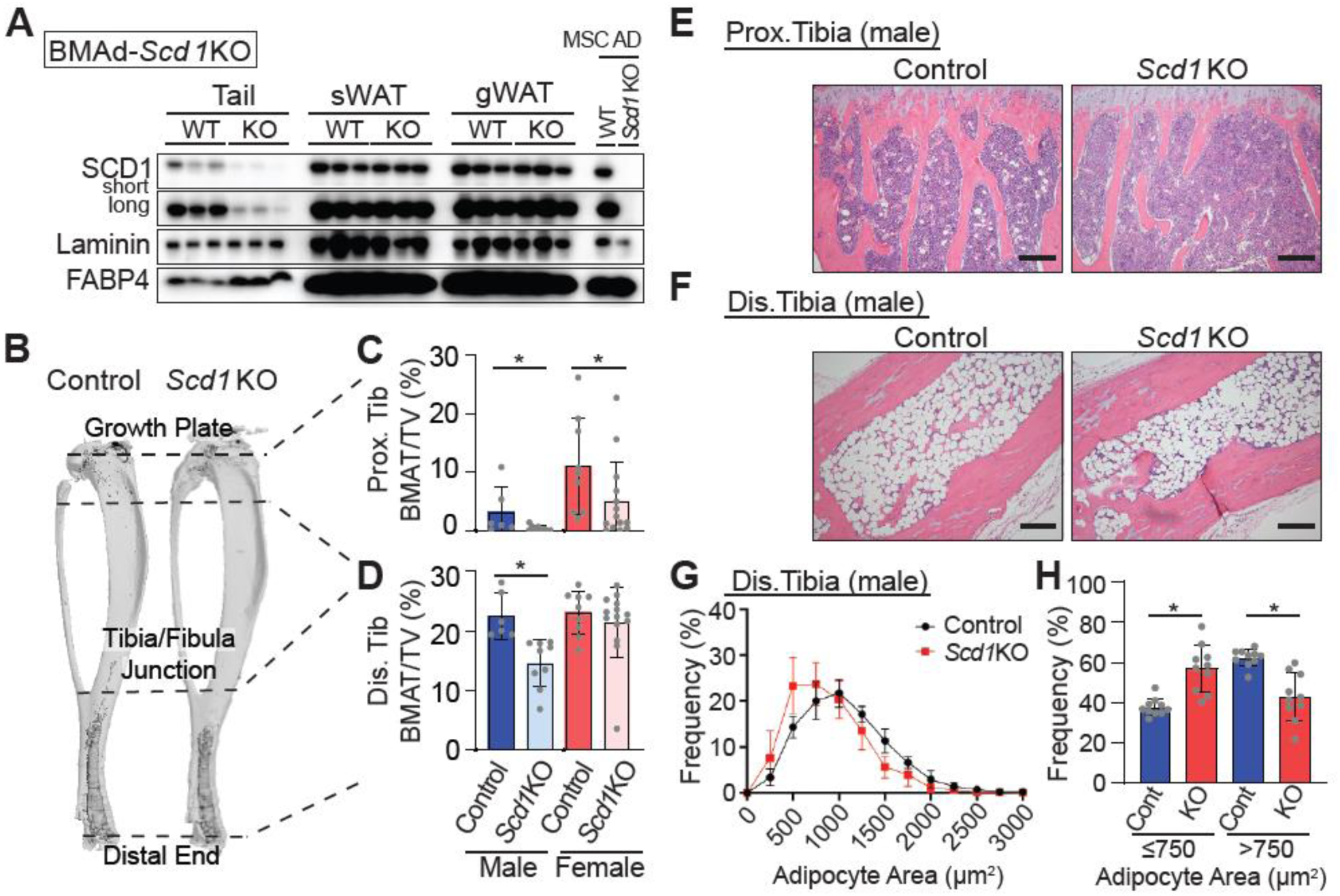
Decreased BMAT in proximal and distal tibiae of *BMAd-Scd1* KO mice. Control and *BMAd-Scd1* KO mice of both sexes were analyzed at 24-26 weeks of age, including 11 male control, 14 male KO, 10 female control, and 13 female KO mice. (**A**) Expression of SCD1 in tail of *Scd1*KO BMAT is reduced but not in sWAT and gWAT depots. Lysates from cultured adipocytes derived from control and *Scd1KO* mice were used as controls. Control and BMAd-*Scd1*KO mice of both sexes were analyzed at 24-26 weeks of age, including 11 male control, 15 male KO, 10 female control, and 14 female KO mice. (**B**-**D**) Decalcified tibiae were treated with osmium tetroxide and quantified by μCT analyses to measure the BMAT volume at proximal and distal ends of the tibia. Data are presented as mean ± SD \**p* < 0.05. (**E** and **F**) Representative images of male proximal tibial BMAT (**E**) and distal tibial BMAT (**F**). Scale bar; 200 μm. (**G** and **H**) Quantitative analyses of BMAd sizes were performed with MetaMorph software. Data are presented as mean ± SD \**p* < 0.05.

### *Scd1*KO adipocytes do not die via canonical apoptotic or necroptotic pathways but undergo autophagy-dependent cell death (ADCD)

While death of cancer cells has been linked to inhibition of SCD1 activity ^22^, mechanisms associated with cell death induced by SCD1 inhibition vary, including apoptosis, ferroptosis, and autophagy ^22^. One potential explanation for this diversity may stem from varying levels of SCD1 expression in cancer cell models (**Figure S6**) and differing extent of SCD1 inhibition ^23–27^. We next investigated which type of cell death occurs in the adipocytes deficient for SCD1. Apoptotic signaling activates a caspase cascade; the initiator caspases in turn cleave and activate downstream effector caspases. However, there was no induction of cleaved caspase 3 in either control or *Scd1*KO adipocytes, whereas adipocytes treated with apoptosis inducers such as thapsigargin or tunicamycin showed robust cleavage of caspase 3 (**Figure S7A**). Necroptosis is a programmed form of necrosis that involves phosphorylation of MLKL. Whereas neither control nor *Scd1*KO adipocyte samples showed evident MLKL phosphorylation, induction of necroptosis was observed in these adipocytes treated with the necroptosis inducers (TSZ: TNF-α, SM-164, and VAD-fmk) (**Figure S7B**). Consistent with these findings, the apoptotic inhibitor ZVAD and three necroptotic inhibitors (i.e. necrostatin, GSK 872, or necrosulfonamide) did not prevent cell death induced by SCD1 deletion in adipocytes (**Figure S7C**). Although SCD1 inhibition has been reported to trigger ferroptosis and death of ovarian cancer cells ^22,27^, inhibitors of ferroptosis (i.e. ferrostatin, liproxstatin 1, or SRS 11-92) were not sufficient to rescue death of *Scd1*KO adipocytes, as compared to supplementation with MUFAs (**Figure S7C** and **S7D**). To confirm the absence of apoptosis characteristics, such as nuclear condensation or apoptotic body formation, and necrosis characteristics, such as cell swelling, we evaluated adipocyte morphology with transmission electron microscopy (TEM). We did not observe the typical indications of apoptosis, such as chromatin condensation and fragmentation, or the main features of necrosis, such as cell swelling (**Figure 5A**). Importantly, unlike control adipocytes, which display uniformly electron-dense lipid droplets, *Scd1*KO adipocytes contain many vacuoles with differing electron densities (**Figure 5A**), and likely correspond to brown structures observed with light microscopy (**Figure 2B** and **3A**).

**Figure 5.**
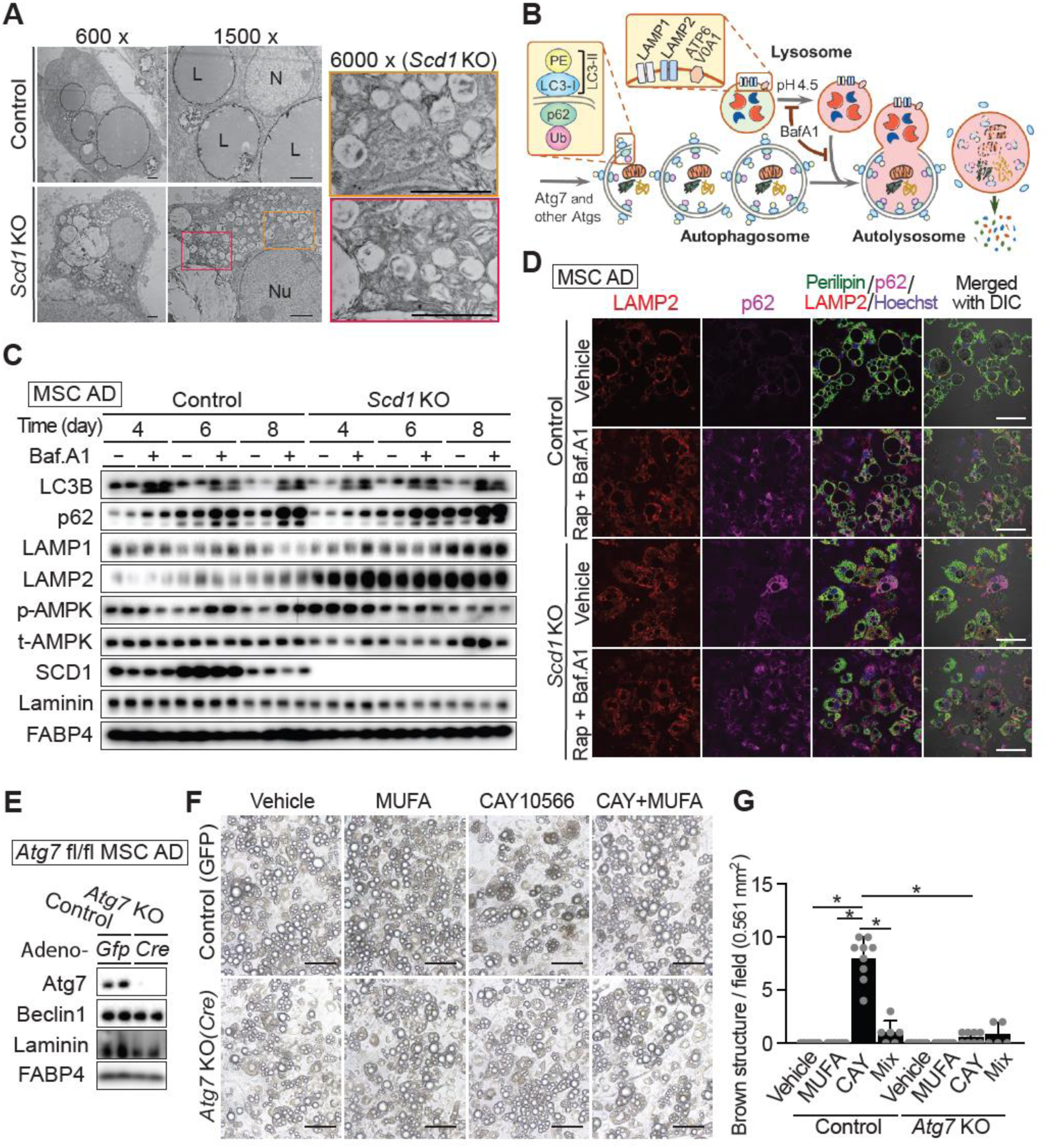
*Scd1*KO adipocytes undergo ADCD rather than by canonical apoptotic or necroptotic pathways. (**A**) MSC adipocytes were fixed on day 8 after differentiation for examination by transmission electron microscopy (TEM). Scale bar; 2 μm (**B**) Schematic diagram of different steps of autophagy. (**C**) To prevent autophagy induced by nutrient deprivation, adipocytes from MSCs at indicated times after differentiation were changed with fresh media one day before assay, then with fresh DMEM/F12 (7.8 mM glucose) with 2% FBS for 4 hrs before harvesting with or without 100 nM of bafilomycin A for 2 hrs. Autophagy-related proteins in control and *Scd1*KO adipocytes were examined by immunoblotting using indicated primary antibodies. (**D**) Immunocytochemistry to detect lysosome-associated membrane protein-2 (LAMP2; red), p62 as an autophagic marker (deep red), perilipin (lipid droplet; green), or Hoechst 33342 (nuclei; blue) on MSC adipocytes after treatment with 100 nM bafilomycin A and 10 nM rapamycin for 3 hrs. Scale bar indicates 50 μm. (**E**-**G**) Autophagy is required for SCD1 inhibition to cause adipocyte death. MSCs from *Atg7*^fl/fl^ mice were induced to differentiate and at day 4, adipocytes were infected with adenoviral vectors carrying *Gfp* (control) or *Cre* (Atg7KO) genes. (**E**) Immunoblot analysis of ATG7 expression and controls for autophagy (beclin), loading (laminin) and adipogenesis (FABP4). (**F**) Following adenoviral infection, adipocytes were cultured with 30 nM of CAY10566 from Day 6 to Day 10. Treatment with 50 µM each of palmitoleic (C16:1) and oleic (C18:1) acids was used as the standard for rescue of cell viability. Scale bar indicates 50 μm. (**G**) The average number of brown structures per field (0.561 mm^2^) was quantified after photomicroscopy of 6 fields per group. \**p* < 0.05. Data shown in (**C**)-(**F**) are representative of at least 3 independent experiments.

Large numbers of small vacuoles present in the cytoplasm is a characteristic feature of ADCD ^13^. It has been proposed that three criteria must be met to determine whether cells die by ADCD. These are to 1) exclude other forms of cell death, 2) observe an increase in autophagic flux, and 3) determine whether blockade of autophagy rescues cell viability ^13,16^. We did not find evidence for death by apoptosis, necroptosis, or ferroptosis (Figure S7); thus, we evaluated autophagic flux by accumulation of lipidated LC3B (LC3B-II) and p62 (**Figure 5B**) and found no obvious change in basal rate of autophagy between control and *Scd1*KO adipocytes (**Figure 5C**). In addition, inhibition of lysosomal v-ATPase activity with bafilomycin A1 increased LC3-II and p62 similarly between genotypes (**Figure 5C**). Interestingly, lysosomal markers including LAMP1 and LAMP2 were increased in *Scd1*KO adipocytes and with bafilomycin A1 treatment. We speculated that immunoblot using whole cell lysates might not detect increased autophagic flux in our model since *Scd1*KO adipocytes did not undergo synchronized cell death (**Figure 2A**). Consistent with this idea, immunocytochemical analyses of p62, which is recruited to autophagosomes forming punctate structures ^28^, revealed that *Scd1*KO adipocytes had increased p62 puncta at baseline, and following bafilomycin A1 and rapamycin treatment (**Figure 5D**). Finally, to evaluate whether blockage of autophagy caused cell death upon SCD1 deletion, we generated ATG7 knockout adipocytes by infecting adenoviral Cre recombinase into *Atg7*^fl/fl^ MSC adipocytes (**Figure 5E**). Whereas brown structures were significantly increased upon SCD1 inhibition with CAY10566 (30 nM for 4 days) in control GFP-infected adipocytes, *Atg7*KO adipocytes did not form brown structures or undergo cell death with the same treatment (**Figure 5F** and **5G**). Taken together, these data indicate that inhibition of SCD1 in adipocytes leads to ADCD *in vitro*. Of note, although TEM analyses showed more vacuole formation in *Scd1*KO BMAds compared to control BMAds (**Figure S7E**), we were unable to confirm whether these are equivalent to vacuoles observed in *Scd1*KO adipocytes *in vitro*.

### Impaired lysosomal acidification is necessary for death of *Scd1*KO adipocytes

Increased expression of lysosomal proteins in *Scd1*KO adipocytes (**Figure 5C**) led us to investigate further the potential relationships between lysosomal function and *Scd1*KO cell death. Lysosome isolation and enrichment from both control and *Scd1*KO adipocytes were achieved by differential centrifugation and density gradient fractionation. Exclusion of basement membrane after separation is shown by the absence of laminin **(Figure 6A)**. Surprisingly, immunoblot analyses revealed reduced expression of LAMP1 and LAMP2 proteins in *Scd1*KO lysosomes whereas their abundance in whole-cell lysate was elevated (**Figure 6A**). Expression of V-ATPase (ATP6V0A1), which pumps protons into the lysosome, is also decreased in *Scd1*KO lysosomes relative to expression of a late endosomal/lysosomal protein, ABHD6 (**Figure 6A**). To identify and quantify abundance of purified proteins, we performed proteomics of lysosomal enrichments from both genotypes. Gene Set Enrichment Analysis (GSEA) of lysosomal proteomics data was conducted using the Gene Ontology gene set with plots separated into GO Biological Process (GOPB). Upregulated pathways in the *Scd1*KO lysosomes were related to mitochondria (**Figure 6B**), perhaps indicative of dysregulated fusion of mitophagic vacuoles with lysosomes, or increased tethering of lysosomes with mitochondria. ^29^ In contrast, GOBP predicts downregulated pathways related to lysosome including Protein Localization to Lysosome, and Lytic Vacuole Transport (**Figure 6B**).

**Figure 6.**
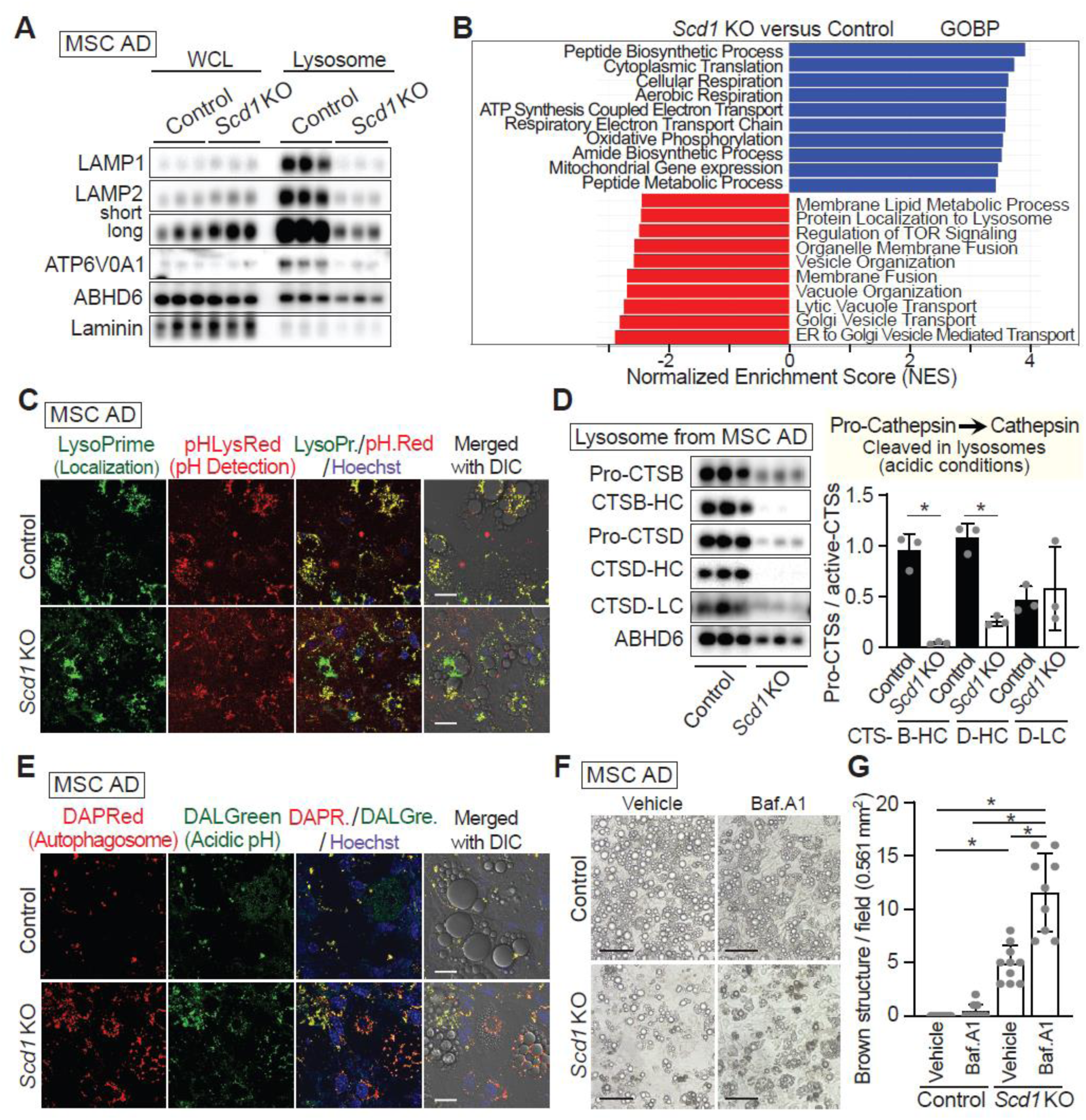
Impaired lysosomal acidification in *Scd1*KO adipocytes is necessary for death of *Scd1*KO adipocytes. (**A** and **B**) Lysosome isolation and enrichment for Day 6 control and *Scd1*KO adipocytes was achieved by cell disruption followed by differential centrifugation and density gradient fractionation. (**A**) Whole cell lysate (WCL) and lysosomal lysate from control or *Scd1*KO adipocytes were examined by immunoblotting using primary antibodies to lysosome markers: LAMP1/2, ATP6V0A1, and ABHD6: Alpha/Beta-Hydrolase Domain-Containing 6. (**B**) Gene Set Enrichment Analysis (GSEA) of lysosomal proteomics data (*n* = 4) was conducted using the Gene Ontology gene set, with plots separated into GO Biological Process (GOBP). The visualizations display the top 10 upregulated and top 10 downregulated pathways, all of which have FDR-adjusted *p*-values < 0.05. (**C**) Reduced numbers of intact lysosomes in *Scd1*KO adipocytes. MSC adipocytes were stained with LysoPrime Green for specific localization of lysosomes, and pHLys Red as a measure of lysosomal pH. Scale bar indicates 20 μm. (**D**) Impaired conversion of pro-cathepsins to active cathepsins in *Scd1*KO lysosomes. Immunoblotting of purified lysosomal fractions with to detect pro-cathepsins and cathepsins. Pro-cathepsin and cathepsin ratio quantified by ImageJ. *n* = 3, data are mean ± SD. \**p* < 0.05. CTSB: cathepsin B, CTSD: cathepsin D (**E**) Decreased autolysosomes in *Scd1*KO adipocytes. Cultured adipocytes were stained with DAPRed for autophagosome detection, and DALGreen as a measure of vesicle acidity. Scale bar indicates 20 μm. (**F**-**G**) Lysosomal acidification in *Scd1*KO adipocytes is necessary for *Scd1*KO cell death. Both control and *Scd1*KO MSC adipocytes were treated with 30 nM bafilomycin A1 from Day 4 after differentiation for 24 hrs, then photos were taken at Day 6. Scale bar indicates 50 μm. (**F**). Brown structures per field (0.561 mm^2^) were quantified in 10 different fields/ in each group (**G**). \**p* < 0.05.

As noted above, we observed decreased recovery of proteins that regulate lysosomal pH including LAMP1/2 and ATP6V0A1(V-ATPase) (**Figure 6A**) ^30^; thus, we hypothesized that lysosomes in *Scd1*KO adipocytes have impaired acidification. To test this hypothesis, MSC adipocytes were stained with LysoPrime Green, a lysosome probe resistant to pH change, for localization of lysosomes, and pHLysRed, a specific probe for acidic lysosome, to indicate extent of lysosomal acidity. Whereas acidic punctate lysosomal structures were readily observed in control adipocytes as indicated by yellow dots in the merged picture (**Figure 6C** top), a subset of *Scd1*KO adipocytes were unable to acidify lysosomal vacuoles, and thus were green due to staining by LysoPrime but not by pHLysRed (**Figure 6C** bottom). To confirm that *Scd1*KO adipocytes have impaired lysosomal acidification, we measured conversion of pro-cathepsin to cathepsin, which occurs in acidic lysosomes. Whereas pro-cathepsin B/D and active cathepsins were observed in a ∼1:1 ratio in control adipocytes, very little active form of cathepsins was observed in *Scd1*KO lysosomes (**Figure 6D**), suggesting that lysosomal acidification was indeed impaired in *Scd1*KO adipocytes.

We next investigated autolysosome formation with DAPRed for detecting autophagosomes, and vacuole acidity with DALGreen, whose fluorescent becomes stronger as acidity increases. In control adipocytes, autolysosomes were observed as yellow dots in the merged picture, suggesting that autophagosomes were efficiently fused with lysosomes (**Figure 6E**). On the other hand, two distinct staining patterns were observed in *Scd1*KO adipocytes. Firstly, there were adipocytes with numerous red autophagosome puncta that were not acidic (**Figure 6E**), suggesting that SCD1 deficiency might cause impaired fusion between autophagosomes and lysosomes, or that autophagosomes might have fused successfully with non-acidic lysosomes. Secondly, there were adipocytes with a large number of acidified autolysosomes, as indicated by the yellow color in the merged picture (**Figure 6E**). Aside from the fact that *Scd1*KO adipocytes did not undergo synchronized cell death (**Figure 2A**), the steady state degree of autophagic compensation and decompensation within *Scd1*KO adipocytes did not result in elevated autophagy flux, as detected by immunoblotting. However, prior to death, *Scd1*KO adipocytes have an excess of autophagosomes, which may result from an imbalance between autophagy initiation and lysosomal degradation.

To evaluate importance of lysosomal acidification on death of *Scd1*KO adipocytes, control and *Scd1*KO adipocytes were treated with bafilomycin A1, from Day four of differentiation for 24 hrs. At day six of adipogenesis, the number of brown structures was increased in *Scd1*KO adipocytes treated with bafilomycin A1 compared with vehicle, and also when compared to control cells treated with bafilomycin A1 (**Figure 6F** and **6G**). Taken together, these data indicate that SCD1 is required to maintain lysosomal acidity, and that impairment of this process results in accumulation of autophagosomes and death of *Scd1*KO adipocytes.

## DISCUSSION

### Comparison of SCD1 knockout phenotypes between our study and others

The genetic basis for a spontaneous mutant mouse, Asebia (ab), characterized by hypoplastic sebaceous glands and alopecia, was determined by positional cloning to be a deletion in *Scd1* ^31,32^. The conclusion that the *Scd1* gene mutation is responsible for the asebia phenotype was further supported by global *Scd1* gene knockout studies ^5^. Whereas global deletion of *Scd1* protected mice against obesity induced by a high carbohydrate diet (HCD), and high-fat diet (HFD), resistance to adiposity was chiefly attributed to increased metabolism. This phenotype was not replicated in tissue-specific KO mice in which SCD1 is highly expressed, such as in the liver ^6^, adipose tissue ^7^, or even in a double KO of *Scd1* in both liver and adipose tissue ^8^. Only skin-specific SCD1 knockout mice increased metabolism to maintain body temperature to compensate for loss of skin-insulating factors ^4,9^.

A recent report demonstrated that *Scd1* deficiency in adipose tissue-derived mesenchymal stem cells promotes *de novo* beige adipogenesis ^19^. The observation that loss of *Scd1* potentiates beige adipocyte formation came from studies on *Scd1*^ab-Xyk^ mice, which exhibit sebaceous gland abnormalities due to SCD1 function loss ^33^. These mice presumably display a hypermetabolic lean phenotype due to the loss of skin-insulating factors, similar to those seen in global *Scd1*KO (GKO) mice ^5^ and *Scd1*^ab-2J^ mice ^31,32^. This effect, including adipose beiging, was particularly marked when the mice were housed at 22°C, a temperature below their thermoneutral point ^3^. In our study, we did not observe that beige adipocyte markers were induced in *Scd1*KO adipocytes derived from MSCs. This finding was supported in *Scd1*KO adipocytes originating from *Scd1*^floxed^ stromal-vascular cells infected with the *Cre* virus. Divergence in our findings on beige adipocyte formation likely arises from differences in experimental models studied. Of importance, we identified the induction of brown structures and cell death in several mouse adipocyte models of SCD1-deficiency. This particular phenomenon was also evident in adipocytes derived from human stromal-vascular cells treated with the SCD1 inhibitor. Additionally, brown-black dots appear in a few representative pictures of *Scd1*^tm1Ntam^ adipocytes ^19^, although it is unknown whether a subset of these structures represents features of cell death. Taken together, these findings suggest that inhibition of SCD1 broadly triggers formation of brown structures and death of adipocytes *in vitro*.

In addition to the hypoplastic sebaceous glands ^5^, pancreas organogenesis in global *Scd1* knockout mice also exhibited decreased formation of β-cells, coinciding with reduced expression of β-cell signature genes, including Pdx1, MafA, and Neurod1 ^34^. It would be intriguing to investigate whether these phenotypes are related to cell death caused by the lack of endogenous MUFAs due to SCD1 deletion. There is no documented research pertaining to cell death in liver-, adipose-, and double liver/adipose SCD1 KO mice ^6–8^. One explanation for this reason is that targeted cells are unlikely to completely depend on endogenous MUFA production and might instead utilize MUFAs taken in from circulation. For example, hepatocytes of liver *Scd1* KO mice obtain exogenous MUFA sources from diet and adipose tissues. If *Scd1*KO hepatocytes use circulating MUFAs to meet their needs, the proposed cell death may not occur, even without *de novo* MUFA synthesis - a hypothesis supported by our observation that MUFA supplementation rescued *Scd1*KO adipocytes from cell death. However, high-level oleate supplementation as triolein did not rescue the sebaceous gland phenotype in the global *Scd1* KO or increase wax levels. This suggests that sebaceous glands depend more on endogenous MUFA generation by SCD1 rather than dietary MUFA incorporation, or that introduction of MUFA supplements was too late to reverse sebaceous cell death processes. Homozygous asebic mice display impaired hair growth at only seven days of age, supporting the latter hypothesis ^32^. This dependence on *de novo* MUFA synthesis also appears to be true for rBMAT and male cBMAT, since reduced numbers of BMAds were observed in mice lacking SCD1 in BMAT. Although targeted cells that largely rely on endogenous MUFAs might die due to insufficient MUFAs following SCD1 deletion, any decrease in cell numbers might not be visible due to asynchronous cell death, and development of new cells. Mouse liver, for example, typically begins to regenerate within hours after injury and can restore lost liver mass within approximately 5-7 days ^35^. Similarly, adipocytes in mice regenerate within 4-6 weeks after cell death ^36^.

### Scd1KO adipocytes undergoes ADCD

There are numerous reports linking SCD1 to cancer cell death ^22^, yet the specific cell death pathways involved - including apoptosis, ferroptosis, and autophagy - remain controversial. One potential explanation for this diversity of pathways might stem from varying SCD1 expression in different cells, as well as differential concentrations of SCD1 inhibitors used in various studies. It is well established that when cells are unable to activate their primary cell death program, they may initiate another one as a compensatory mechanism. Autophagy is primarily recognized for its cytoprotective role in cellular processes. Compelling evidence suggests that inhibition of essential *Atg* genes can accelerate cell death when the target of selective autophagic degradation comprises pro-death proteins, including caspase 8 ^37^ and RIPK1 ^38^. However, autophagy also has the potential to induce cell death particularly during development and in response to stress, a phenomenon classified as ADCD. To conclusively determine the occurrence of ADCD, three criteria were proposed ^13,16^. These include: 1) the exclusion of other forms of cell death, 2) evidence of an increase in autophagic flux, and 3) proof that inhibiting autophagy can rescue the cell from death. Although we weren’t able to exclude all 12 major cell death modes ^39^, we successfully excluded necrosis, apoptosis, and ferroptosis, using TEM and various inhibitors. Despite not detecting increased autophagic flux via immunoblotting, p62 immunocytochemistry revealed that some *Scd1*KO adipocytes were filled with p62 puncta, which further increased with bafilomycin A1 and rapamycin treatment. This observation was supported by the DAPRed (autophagosome) and DALGreen (acidity) stains that revealed the presence of many autolysosomes in *Scd1*KO adipocytes. Another distinct group of adipocytes demonstrated numerous autophagosome puncta that lacked acidification, suggesting potential impairment in the fusion between autophagosomes and lysosomes. This might lead to cell death, possibly representing a stage of decompensation when a lysosome cannot manage an enhanced autophagosome. This dynamic, consisting of compensational and decompensation phases, resulted in little change in autophagic flux as detected by immunoblotting in *Scd1*KO adipocytes. However, ATG7 knockout adipocytes were resistant to cell death induced by SCD1 inhibition. Taken together, these findings suggest that inhibition of SCD1 in adipocytes induces ADCD *in vitro*.

Morphologically, three types of cell death, including ADCD, are associated with multiple vacuole formations ^40^. We excluded paraptosis, characterized by cytoplasmic vacuolation that involves the swelling of the endoplasmic reticulum and mitochondria, as no mitochondrial swelling was observed in *Scd1*KO adipocytes. In addition, methuosis is characterized by electron-lucent cytoplasmic fluid-filled vacuoles resulting from macropinocytosis ^41^, which also does not fully align with our observations. We observed vacuoles in *Scd1*KO adipocytes by TEM whose contents appeared to have different cargo from controls, or appeared empty, consistent with the characteristics of ADCD due to increased accumulation of autophagosomes positive for p62 positive puncta and lacking acidification.

### Critical roles of lipogenesis for autophagy

Although the exact source of the autophagosomal membrane is still a debated topic, studies suggest that autophagosomes might originate from the ER or from ER-related membranes ^15^. MUFAs, which are generated by the ER-associated enzyme SCD1, are key components in the biosynthesis of lipids including phospholipids. Since deletion of SCD1 shifts lipid composition from species containing monounsaturated lipids toward species containing saturated lipids in the lipidomic data from whole adipocytes, quantity of MUFAs incorporated into the autophagosome membrane could potentially influence the efficiency and specificity of autophagic processes. Although existing studies provide evidence that lipids are involved in controlling autophagy, they demonstrate a relationship between the composition of phospholipids, the storage of lipid droplets, ER stress, and the biogenesis of autophagosomes ^42–44^.

There are two recent, intriguing papers related to autophagosome membrane assembly. The yeast Acyl-CoA synthetase, Fatty acid activation protein 1 (Faa1), channels fatty acids into *de novo* phospholipid synthesis and promotes assembly of newly synthesized phospholipids into autophagic membranes ^45^. Another paper reported that FASN colocalizes with nascent autophagosomes ^17^. A deficiency of FASN in adipocytes, both *in vitro* and *in vivo*, markedly impairs autophagy, causing substantial accumulation of autophagosomes ^17^. There is no further information regarding the fate of *Fasn*KO adipocytes that have accumulated autophagosomes, including cell death. Interference with cellular transport due to packed autophagosomes, or the disappearance of intact organelles without being sequestered by an autophagosome, are potential mechanisms of ADCD, although the precise mechanisms remain unclear ^13,14^. Thus, it is possible that cells with a massive accumulation of autophagosomes in *Fasn*KO adipocytes might also undergo cell death. The results from *Fasn*KO adipocytes uncovered the role of adipocyte DNL in autophagy, as opposed to its relatively minor contribution to adipocyte triacylglycerol storage ^17^. Although palmitate was insufficient to restore autophagic flux in *Fasn*KO adipocytes, we speculate that direct supplementation of MUFAs may restore autophagy activity in *Fasn* KO adipocytes. Our experiments indicate that exogenous palmitate is not an efficient substrate for SCD1 in adipocytes, and that supplementation with MUFAs rescues adipocytes from cell death induced by SCD1 depletion. Our studies propose that the MUFAs produced by SCD1 and potentially incorporated into autophagosome membranes, can influence efficiency and specificity of autophagic processes. Further study is required to define the molecular mechanisms by which MUFAs within the autophagosome membrane regulate autophagy especially in adipocytes.

Several studies address the role of SCD1 or MUFA on autophagosome formation, and autophagosome-lysosome fusion ^11,46–49^. Although the observed phenomena are somewhat similar to those highlighted herein, the conclusions drawn are different. For instance, INS-1E pancreatic β-cells treated with 2 μM A939572, an SCD1 inhibitor, have impairments in autophagic clearance due to defects in autophagosome-lysosome fusion, resulting in apoptosis rather than ADCD, presumably due to ER dysfunction ^46^. Human HCC cells treated with 10 μM CAY10566, another SCD1 inhibitor, induced both apoptosis and autophagy ^49^. In contrast, SCD1 inhibitor treatment of mouse embryonic fibroblasts suppressed autophagy at the earliest stage of autophagosome formation ^11^. Interestingly, treatment of human osteosarcoma U2OS cells with 500 μM oleate induced a non-canonical autophagic response, presenting vacuolar structures colocalizing with the Golgi apparatus ^48^. Potential explanations for these varied results, diverging from ours, might stem from different levels of SCD1 expression across cell types, varied concentrations of SCD1 inhibitors used, differential toxicity of free fatty acids among cell types evaluated, and differences in underlying metabolism of adipocytes and other cell types.

### Scd1KO does not undergo lysosome-mediated cell death

Lysosome-dependent cell death is defined as a cell death event characterized by massive lysosomal leakage and the ectopic action of lysosomal enzymes ^50^. Different forms of stress have been reported to induce lysosomal membrane permeabilization, resulting in lysosomal leakage, which causes an increase in cytosolic acidity and the uncontrolled breakdown of cell components, inducing necrosis ^50^. Lysosomal swelling caused by membrane permeabilization is the key morphological feature of this type of cell death. However, we did not observe evidence of lysosome-dependent cell death in our immunocytochemistry with lysosomal markers, staining with dyes for lysosomes, or TEM.

Taken together, our results demonstrate that cool adaptation of adipocytes results in elevated expression of SCD1 in cultured adipocytes and in adipose depots. Our exploration of the functionality of this observation demonstrated that *in vitro* inhibition of SCD1 in adipocytes leads to ADCD, and *in vivo* depletion leads to loss of BMAds. *De novo* synthesis of MUFAs by SCD1 is required for fusion of autophagosomes to lysosomes, highlighting essential roles of SCD1 in proper autophagic processes and cell viability. Furthermore, MUFAs play essential roles in maintaining lysosomal and autolysosomal acidification in adipocytes. Lack of de novo synthesis of MUFAs by SCD1 results in aberrant accumulation of vacuoles, eventually leading to cell death.

## Method Details

### Animals

*Scd1*^fl/fl^ mice were provided by Dr. James Ntambi (University of Wisconsin-Madison) and were crossed with BMAd-specific *Cre* mice ^20^ to generate mice with *Scd1* deficiency in bone marrow adipocytes. All animal studies were performed in compliance with policies of the University of Michigan Institutional Animal Care and Use Committee. The protocol number is PRO00009687. All animals were housed in a 12-h light/12-h dark cycle with free access to food and water. Body composition: lean, fat masses were measured by EchoMRI-100H.

### Glucose and insulin tolerance tests

For glucose tolerance tests, mice were fasted for 16 h and then given glucose (1 mg/kg body weight) via intraperitoneal injection. For insulin tolerance tests, mice were fasted for 6 h and then intraperitoneally injected 0.5 U/kg body weight of insulin (Eli Lilly). Glucose concentrations were monitored in blood collected from the tail vein at 0, 15, 30, 60, and 120 min after injection using a glucometer and Contour Next blood glucose strips (Bayer AG).

### Adipocyte and stromal-vascular cell fractionation

Posterior subcutaneous WAT (sWAT) was excised from mice as previously described ^3^. WAT depots from *Scd1*^fl/fl^ mice were minced with scissors and digested for 1 h at 37°C in 2 mg/ml collagenase type I (Worthington Biochemical) in Krebs-Ringer-HEPES (KRH; pH 7.4) buffer containing 3% fatty acid-free bovine serum albumin (BSA; Gold Biotechnology, St. Louis, NJ), 1 g/L glucose, and 500 nM adenosine. The resulting cell suspensions were filtered through 100 μm cell strainers and centrifuged at 100 x g for 8 min to pellet stromal-vascular cells (SVC) and float buoyant adipocytes. To induce gene recombination in adipocytes, *Scd1*^fl/fl^ stromal vascular cells were treated with adenoviral GFP or adenoviral Cre recombinase (1 × 10^10^ viral particles/ml) in serum-free DMEM:F12 from days 4-6 of differentiation. Adipocytes were then analyzed on day 13 of differentiation. Adenoviruses were obtained from the University of Michigan Vector Core.

### Isolation and differentiation of adipocyte precursors

Mesenchymal precursors were isolated from ears of mice of the indicated genotypes, as previously described ^51^. Cells were maintained in 5% CO_2_ and DMEM/F12 1:1 media (Gibco; Invitrogen) supplemented with 10% FBS (Cytiva HyClone), primocin (InVivoGen), and 10 ng/ml recombinant bFGF (PeproTech). For induction of adipogenesis, recombinant bFGF was removed and replaced with 10% FBS containing 0.5 mM methylisobutylxanthine, 1 μM dexamethasone, 5 μg/ml insulin, and 5 μM rosiglitazone. On day 2, cells were fed 5 μg/ml insulin plus 5 μM troglitazone. On day 4 and every 2 days thereafter, cells were fed with 10% FBS-supplemented media. All the media conditions used in the experiments within this study contain 10% FBS, unless otherwise specified in the individual figure legends. When using 2% FBS, Insulin-Transferrin-Selenium (ITS) (Gibco; Invitrogen) was supplemented.

### Lysosome enrichment from cultured MSC adipocytes

Lysosomal fraction was enriched using Pierce™ Lysosome Enrichment Kit (Thermo Scientific) according to the manufacturer’s instructions. Briefly, both control and *Scd1* KO adipocytes (Day 6 after differentiation; 10 cm dish) were harvested with 2000 µL of Lysosome Enrichment Reagent A, then homogenized by Dounce tissue grinder after vortex on ice. Lysate was mixed with an equal volume of Lysosome Enrichment Reagent B, then centrifuged at 500 × g for 10 minutes at 4°C to obtain the lysosome-containing supernatant. The buoyant lipid cake was removed after the centrifuge step. After overlaying the supernatant containing 15% OptiPrep Media on top of step-wise density gradients (30%, 27%, 23%, 20% and 17%), samples were ultracentrifuged at 145,000 × g for 2 hours at 4°C. Lysosome bands located in the top 2 mL of the gradient after the end of centrifugation was carefully transferred to new tubes, mixed with 3 volumes of PBS to decrease the concentration of the OptiPrep Media, then centrifuged at 18,000 × g for 30 minutes at 4°to obtain the lysosome pellet.

### Detection of specific proteins by immunoblot

After lysis in 1% NP-40, 120 mM NaCl, 50 mM Tris-HCl (pH 7.4), 50 mM NaF, 2 mM EDTA, 1× protease inhibitor cocktail (Sigma-Aldrich), protein concentrations of lysates after centrifugation were measured by BCA protein assay (Thermo Fisher Scientific). Lysates were diluted to equal protein concentrations in lysis buffer and then boiled in SDS sample buffer (20 mM Tris; pH 6.8, 2% SDS, 0.01% bromophenol blue, 10% glycerol, 5% 2-mercaptoethanol) and subjected to SDS-PAGE and immunoblotting according to standard techniques.^3^

### Immunocytochemistry (ICC)

MSCs were seeded on sterile 12 mm #1.5 round coverslips in 12 well plates, then differentiated into adipocytes. Adipocytes with coverslips were incubated with 4% paraformaldehyde for fixation, with 0.5% Triton X-100 in TBS for permeabilization. Adipocytes were then incubated with primary antibodies solution in 2% BSA for 1 hour at room temperature (RT) after blocking with 2% BSA, then incubated with fluorescent-dye conjugated secondary antibodies in 2% BSA for 1 hr at RT. The coverslip was placed onto a glass slide with ProLong Gold Antifade (ThermoFisher) after counter-staining with Hoechst 33342 (Cayman Chemical; 5 µg/mL).

### Cell Imaging with fluorescent dye, and time-lapse image

The LIVE/DEAD Viability/Cytotoxicity Kit (Invitrogen: L3224) was used following the manufacturer’s instructions to evaluate cell survival. Cells were stained with 2 µM calcein-AM (green) and 4 µM ethidium homodimer I (red) for 30 min. To assess dead cells, adipocytes were incubated with 0.2 % Trypan Blue Solution (Fisher Scientific) in Phosphate Buffered Saline for 3min, then washed with DMEM/F12 media for taking photos. Hoechst 33342 was used for staining nuclei of either living or fixed cells. Time-lapse imaging was performed using a Cytation 5 automated imager with BioSpa-System (BioTek; University of Michigan Frankel Cardiovascular Regeneration Core Laboratory.

### Histology

Tibiae were first fixed by formalin, then decalcified with 14% EDTA for at least two weeks, followed by additional fixation in 4% paraformaldehyde. Soft tissues were fixed with formalin, and both tibiae and soft tissues were embedded in paraffin blocks, which were sectioned and stained with hematoxylin and eosin. Stained tibiae and soft tissues were then imaged on an Olympus BX51 microscope. Adipocyte cross-sectional areas were measured using MetaMorph software.

### Osmium tetroxide staining and µCT analysis of bone marrow lipid

Specimens placed in a 19 mm diameter specimen holder and scanned using a microCT system (µCT100 Scanco Medical, Bassersdorf, Switzerland). Scan settings were: voxel size 12 µm, 70 kVp, 114 µA, 0.5 mm AL filter, and integration time 500 ms. Analysis was performed using the manufacturer’s evaluation software, and a fixed global threshold of 28% (280 on a grayscale of 0–1000) for cortical bone and 25.5% (255 on a grayscale of 0-1000) for trabecular bone was used to segment bone from non-bone. A 1 mm region of trabecular bone was analyzed immediately below the growth plate and a 1.5 mm region around the midpoint of the tibia and a region from the tibia-fibula junction to the distal end. After the analysis of bone variables, mouse tibiae were decalcified for staining with osmium tetroxide, using a method previously published^20^. Additionally, a lower threshold (300 grey-scale units) was employed for the quantification of proximal tibial rBMAT because the density of osmium staining is lower due to the smaller adipocyte size. A threshold of 400 grey-scale units was used for cBMAT in the distal tibia.

### Statistics

All data are presented as mean ± SD. When comparing 2 groups, significance was determined using 2-tailed Student t test. When comparing multiple experimental groups, an analysis of variance (ANOVA) was followed by post hoc analyses with Dunnett or Sidak test, as appropriate. Differences were considered significant at p < 0.05 and are indicated with asterisks. For metabolomics data analyses, the proportion of labeling at each carbon position was calculated by dividing each species by total sum of peak areas of all labeled positions. The proportion of data is well known to follow a beta distribution. Beta regression model is an extension of the generalized linear model with an assumption that the response variable follows a beta distribution with values in standard unit interval (0,1).

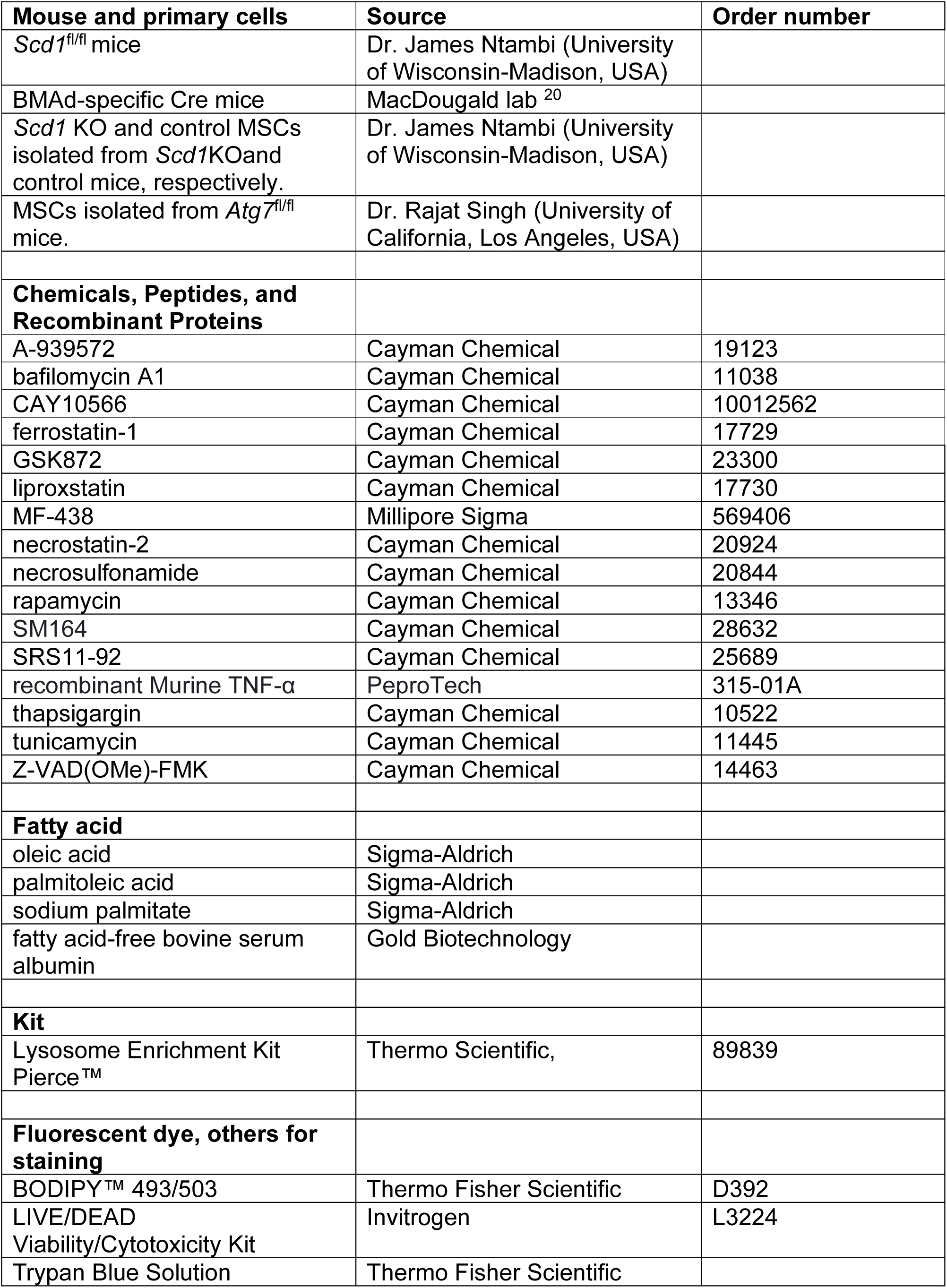
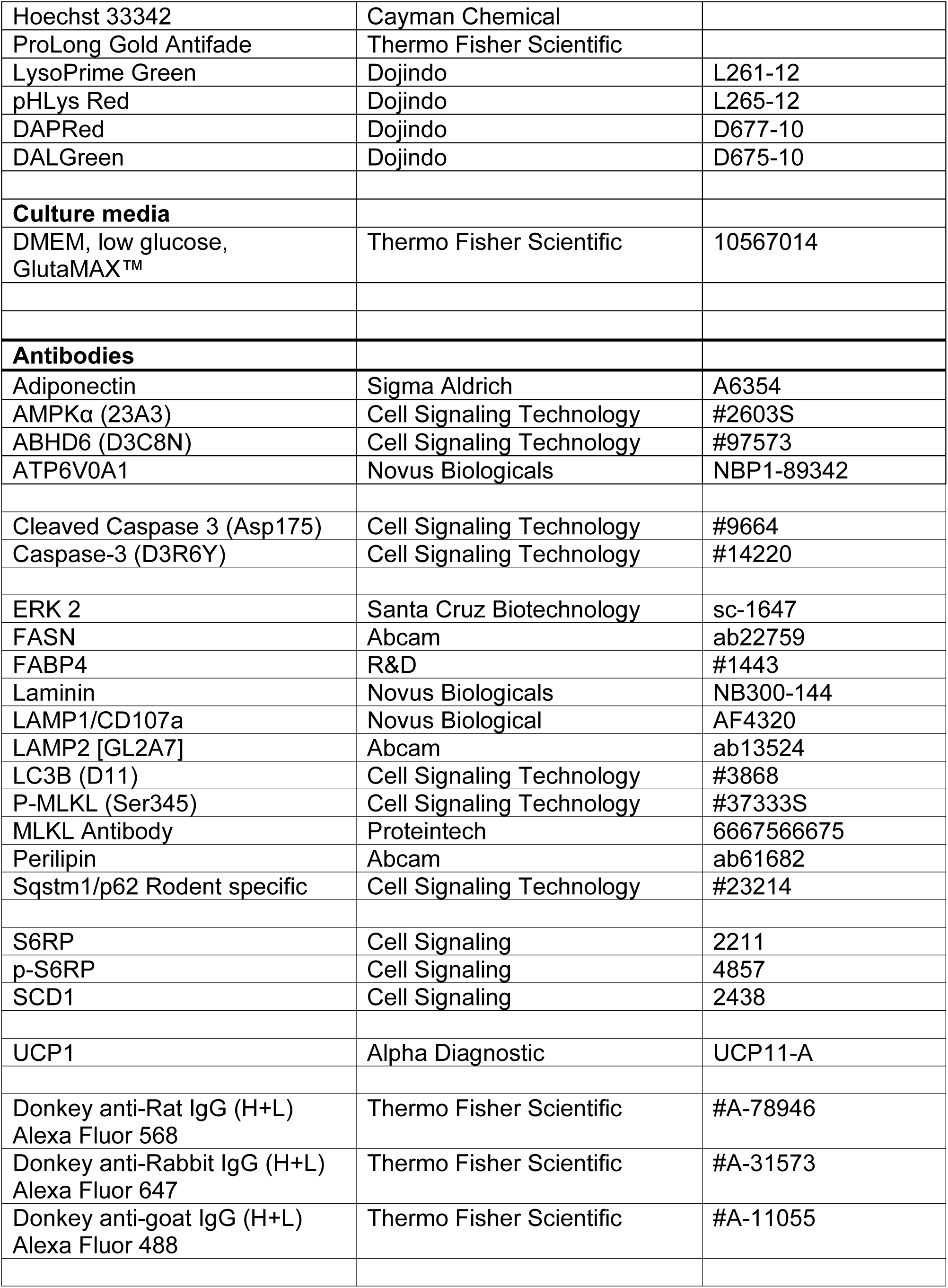

qPCR primers used were as follows;

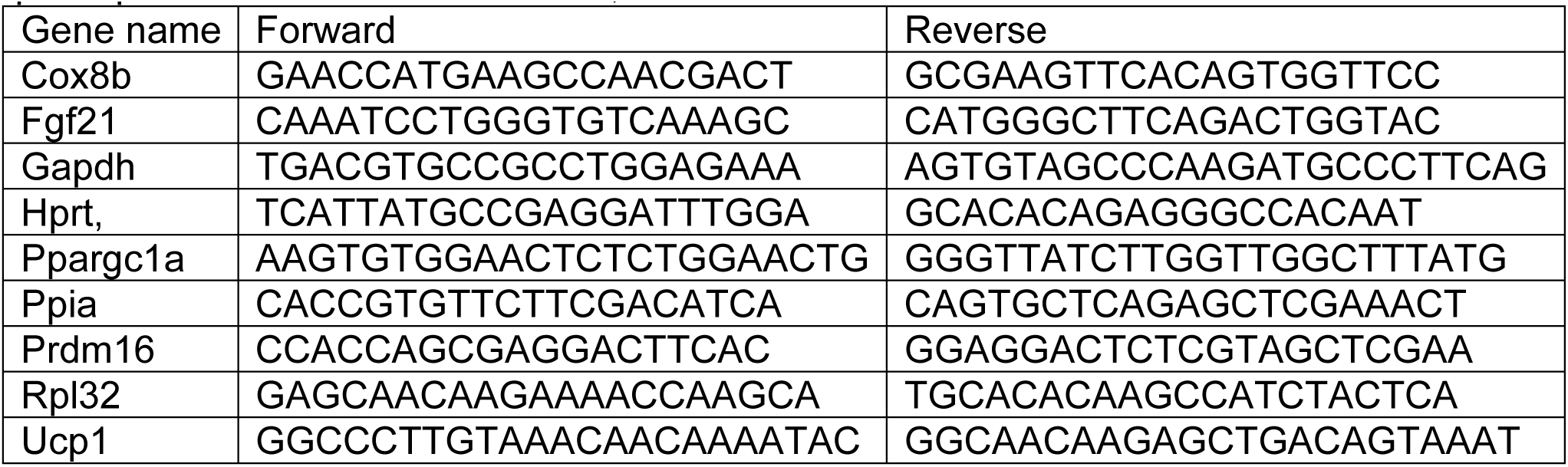

## ACKNOWLEDGEMENTS

This work was supported by grants or fellowships from the NIH to OAM (RO1 DK121759; R01 DK125513, and R01 DK130879), ZL (1P20GM121301), JM (F31 DK135181), KTL (T32 DK071212; F32 DK122654), RLS (T32 DK101357; F32 DK123887), KI (R01DK124709; R01GM145631), and JC (P41 GM108538). ZL was supported by a fellowship from the American Diabetes Association (1-18-PDF-087). This research was also supported by core facilities of the Michigan Integrative Musculoskeletal Health Core Center (P30 AR069620), Michigan Diabetes Research Center (P30 DK020572), Michigan Nutrition and Obesity Center (P30 DK089503), and University of Michigan Frankel Cardiovascular Regeneration Core Laboratory, Histology core (National Institute of Arthritis and Musculoskeletal and Skin Diseases of the National Institutes of Health under P30 AR069620), SoD mCT Core (NIH/NCRR S10RR026475-01). We thank Hannah Thompson, Maria Del Mar Mendez Casillas for technical assistance and members of the MacDougald lab for helpful discussions and assistance.

