## Supplemental Figure 1-7 for "SCD1 and monounsaturated lipids are required for autophagy and survival of adipocytes"

### Supplementary Data

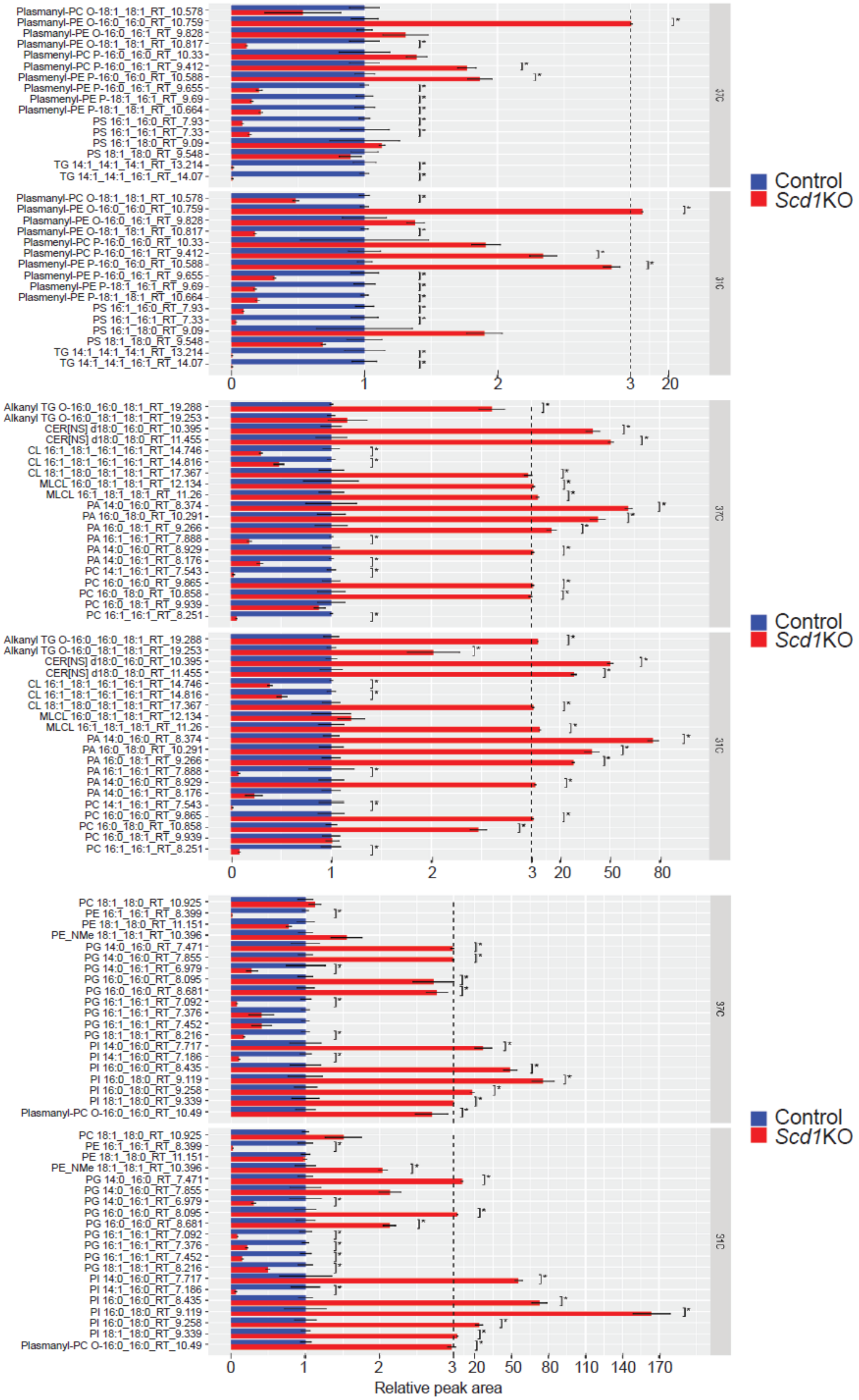
Plasmanyl-PC O-18:1_18:1_RT_10.578
Plasmanyl-PE O-16:0_16:0_RT_10.759
Plasmanyl-PE O-16:0_16:1_RT_9.828
Plasmanyl-PE O-18:1_18:1_RT_10.817
Plasmenyl-PC P-16:0_16:0_RT_10.33
Plasmenyl-PC P-16:0_16:1_RT_9.412
Plasmenyl-PE P-16:0_16:0_RT_10.588
Plasmenyl-PE P-16:0_16:1_RT_9.655
Plasmenyl-PE P-18:1_16:1_RT_9.69
Plasmenyl-PE P-18:1_18:1_RT_10.664
PS 16:1_16:0_RT_7.93
PS 16:1_16:1_RT_7.33
PS 16:1_18:0_RT_9.09
PS 18:1_18:0_RT_9.548
TG 14:1_14:1_14:1_RT_13.214
TG 14:1_14:1_16:1_RT_14.07
37C
31C
0
1
2
3
20
Control
Scd1KO
Alkanyl TG O-16:0_16:0_18:1_RT_19.288
Alkanyl TG O-16:0_18:1_18:1_RT_19.253
CER[NS] d18:0_16:0_RT_10.395
CER[NS] d18:0_18:0_RT_11.455
CL 16:1_18:1_16:1_16:1_RT_14.746
CL 16:1_18:1_16:1_16:1_RT_14.816
CL 18:1_18:0_18:1_18:1_RT_17.367
MLCL 16:0_18:1_18:1_RT_12.134
MLCL 16:1_18:1_18:1_RT_11.26
PA 14:0_16:0_RT_8.374
PA 16:0_18:0_RT_10.291
PA 16:0_18:1_RT_9.266
PA 16:1_16:1_RT_7.888
PA 14:0_16:0_RT_8.929
PA 14:0_16:1_RT_8.176
PC 14:1_16:1_RT_7.543
PC 16:0_16:0_RT_9.865
PC 16:0_18:0_RT_10.858
PC 16:0_18:1_RT_9.939
PC 16:1_16:1_RT_8.251
50
80
PC 18:1_18:0_RT_10.925
PE 16:1_16:1_RT_8.399
PE 18:1_18:0_RT_11.151
PE_NMe 18:1_18:1_RT_10.396
PG 14:0_16:0_RT_7.471
PG 14:0_16:0_RT_7.855
PG 14:0_16:1_RT_6.979
PG 16:0_16:0_RT_8.095
PG 16:0_16:0_RT_8.681
PG 16:1_16:1_RT_7.092
PG 16:1_16:1_RT_7.376
PG 16:1_16:1_RT_7.452
PG 18:1_18:1_RT_8.216
PI 14:0_16:0_RT_7.717
PI 14:1_16:0_RT_7.186
PI 16:0_16:0_RT_8.435
PI 16:0_18:0_RT_9.119
PI 16:0_18:0_RT_9.258
PI 18:1_18:0_RT_9.339
Plasmanyl-PC O-16:0_16:0_RT_10.49
110
140
170
Relative peak area

Figure S1

**Supplemental Figure 1. Effects of cool adaptation and SCD1-deficiency on lipid profile of cultured adipocytes.**

Lipidomic analysis of *Scd1*KO and control adipocytes *in vitro*. Adipocytes were cultured at 37°C or 31°C for 12 days. The peak area for each lipid is expressed as a fold change in *Scd1*KO adipocytes relative to control adipocytes at each temperature (*n* = 3). Log2(Fold Change) values are plotted for lipids containing only 14:0, 16:0, 18:0, 14:1, 16:1, or 18:1.

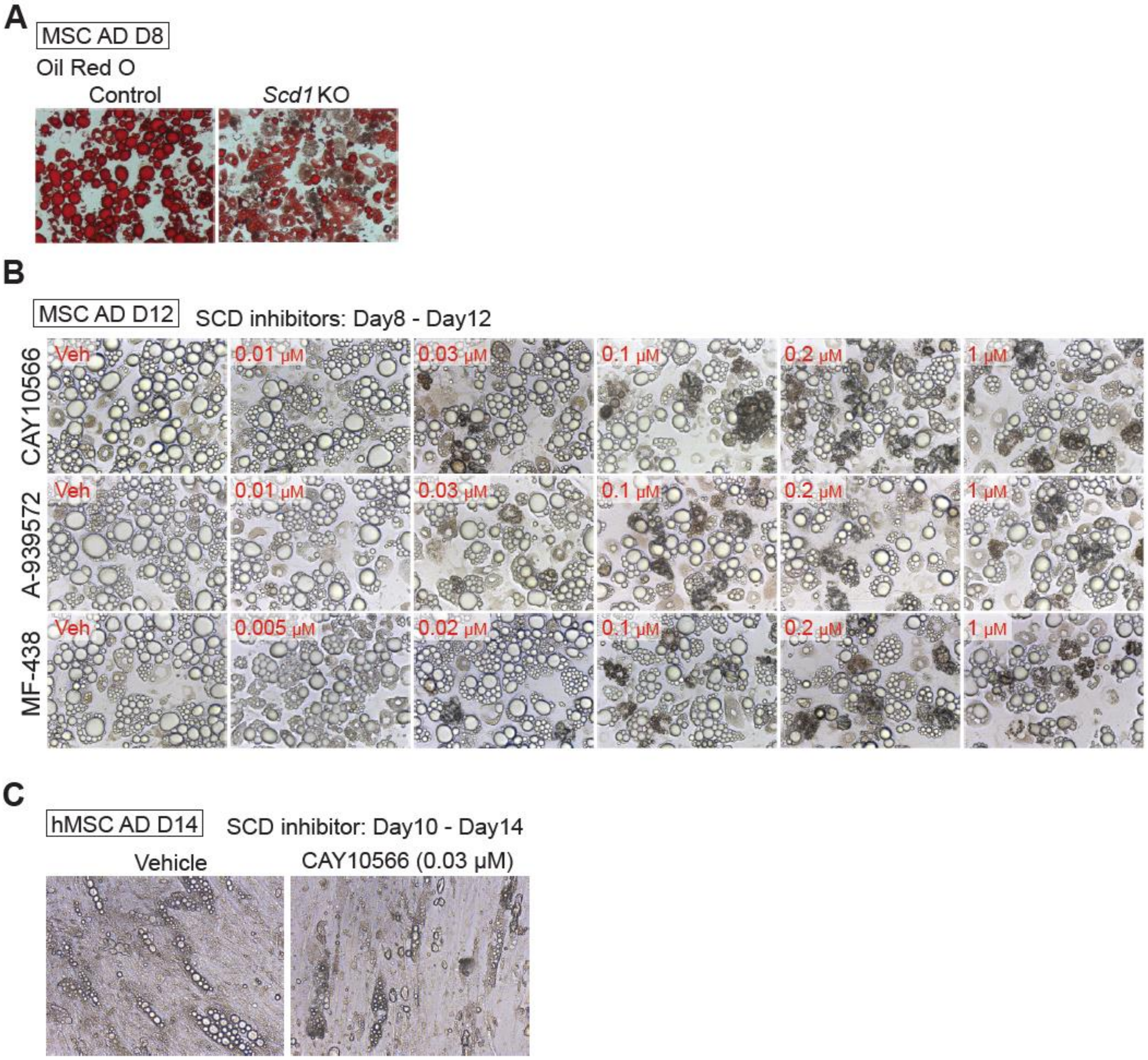


Figure S2

**Supplemental Figure 2. Pharmacological inhibition of SCD1 causes adipocytes to form brown structures.**

(**A**) Photomicrographs of Oil Red-O (ORO) staining in control and *Scd1*KO adipocytes.

(**B**) Phase contrast microscopy of adipocytes were cultured with vehicle or the indicated concentrations of SCD inhibitors from day 8 to 12 of differentiation.

(**C**) Phase contrast microscopy of human precursors treated with 0.03 μM CAY10566 from day 10 to 14 of adipogenesis.

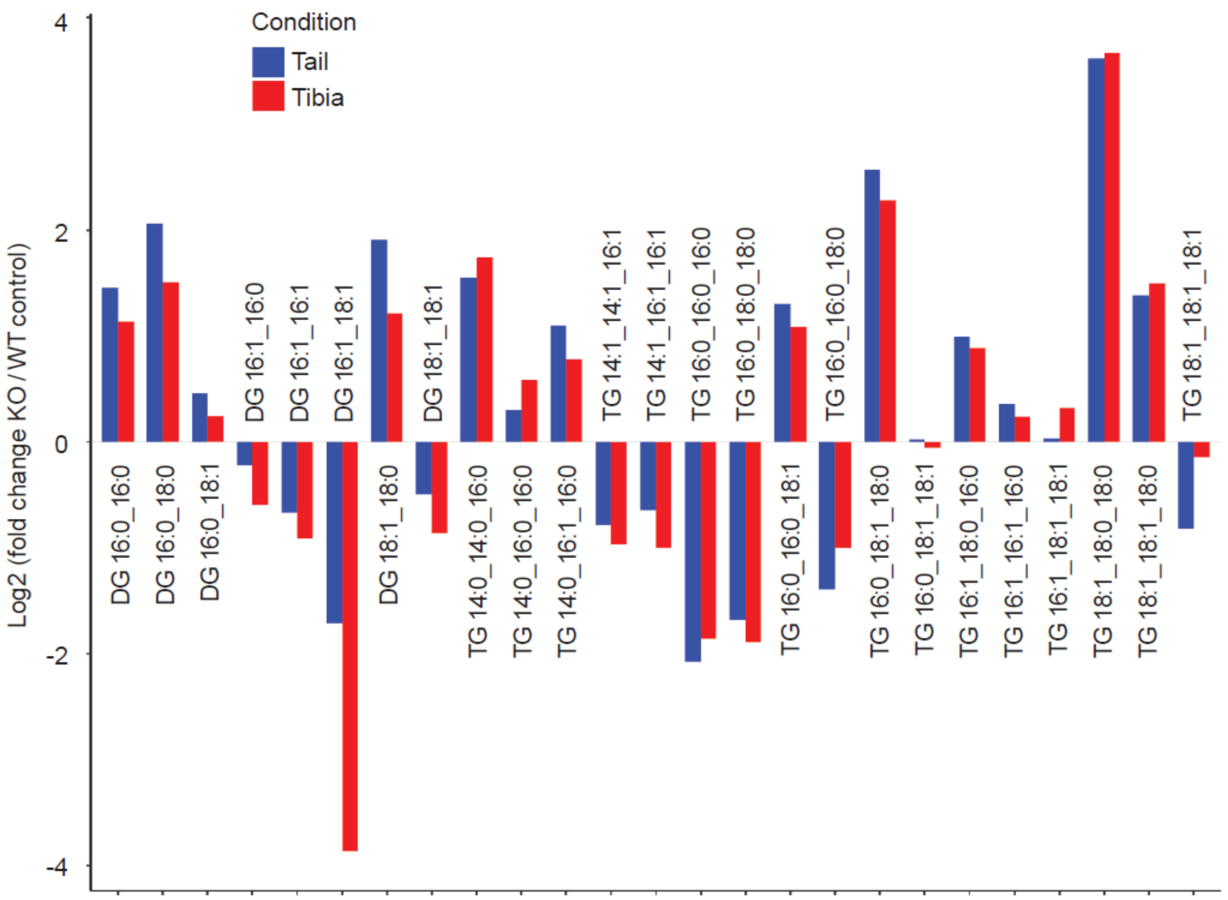


Figure S3

**Supplemental Figure 3**

Lipidomic analysis was conducted on the tibia and distal tibia from female control and BMAd-*Scd1* KO mice, aged 24-26 weeks. These were from the cohort analyzed and reported in Figure 4. The peak area for each lipid is expressed as a fold change in *Scd1*KO tissues relative to control tissues (*n* = 4). Log2 (Fold Change) values are plotted for samples containing exclusively 14:0, 16:0, 18:0, 14:1, 16:1, or 18:1 lipids.

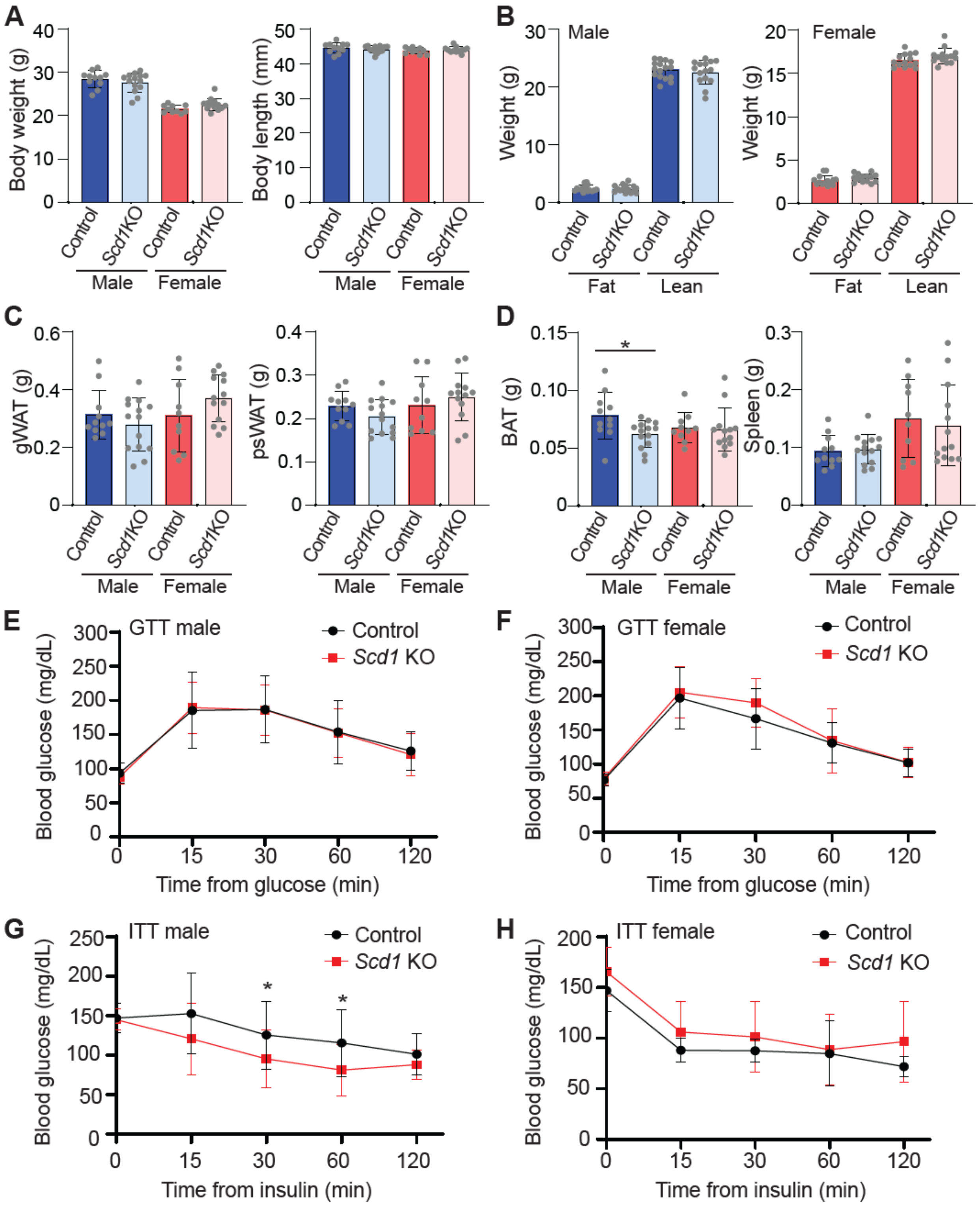
A
Body weight (g)
Body length (mm)
Control
Scd1KO
Male
Female
B
Male
Female
Weight (g)
Fat
Lean
C
gWAT (g)
psWAT (g)
D
BAT (g)
Spleen (g)
E
GTT male
Control
Scd1 KO
Blood glucose (mg/dL)
Time from glucose (min)
F
GTT female
G
ITT male
Time from insulin (min)
H
ITT female

Figure S4

**Supplemental Figure 4. *Scd1*KO mice exhibit phenotypic similarity to control mice in terms of body weight, tissue weights, and glucose tolerance.**

Mice of the indicated genotypes and sexes at 24 - 26 weeks of age were used for investigation. Male (control: $n = 11$, *BMAd-Scd1* KO: $n = 14$) and female (control: $n = 10$, *BMAd-Scd1* KO: $n = 13$).

(**A**) Body weight and length were not changed with genotype in male and female.

(**B**) No genotype-specific differences in lean or fat body masses were observed.

(**C-D**) Weights of gonadal WAT (gWAT), posterior subcutaneous WAT (psWAT)(**C**), brown adipose tissue (BAT), and spleen (**D**). $*p < 0.05$.

(**E**-**F**) No differences in glucose tolerance test (GTT) between genotypes in (**E**) male or (**F**) female mice.

(**G**-**H**) Increased insulin sensitivity in (**G**) male *BMAd-Scd1* KO mice but not (**H**) female mice.

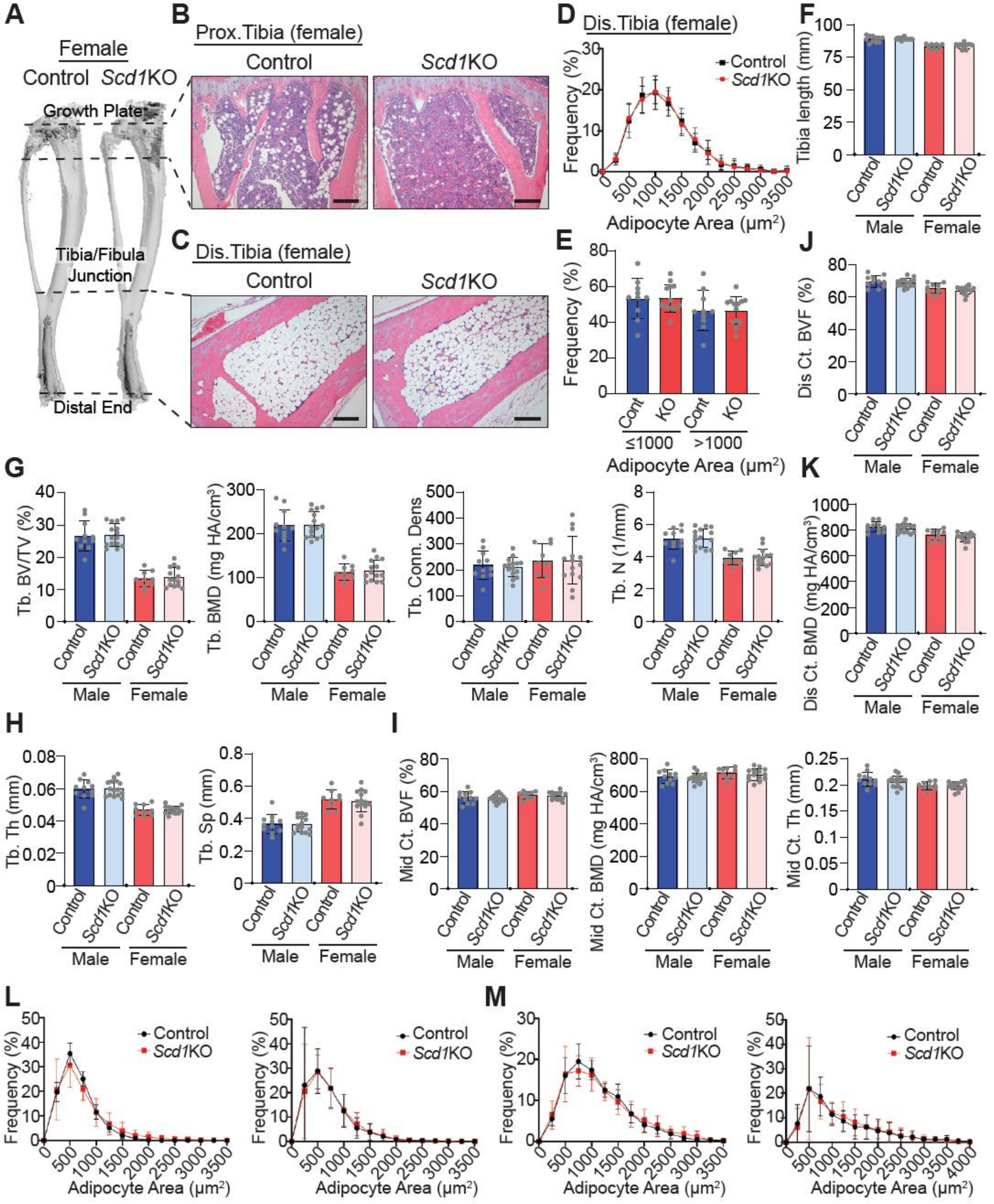
A
Female
Control Scd1KO
Growth Plate
Tibia/Fibula Junction
Distal End
B
Prox.Tibia (female)
Control
Scd1KO
C
Dis.Tibia (female)
Control
Scd1KO
D
Dis.Tibia (female)
Control
Scd1KO
Frequency (%)
Adipocyte Area (μm²)
E
Frequency (%)
Cont
KO
≤1000
>1000
Adipocyte Area (μm²)
F
Tibia length (mm)
Control
Scd1KO
Male
Female
J
Dis Ct. BVF (%)
G
Tb. BV/TV (%)
Tb. BMD (mg HA/cm³)
Tb. Conn. Dens
Tb. N (1/mm)
K
Dis Ct. BMD (mg HA/cm³)
H
Tb. Th (mm)
Tb. Sp (mm)
I
Mid Ct. BVF (%)
Mid Ct. BMD (mg HA/cm³)
Mid Ct. Th (mm)
L
M
Frequency (%)
Control
Scd1KO
Adipocyte Area (μm²)

Figure S5

**Supplemental Figure 5. Deficiency of SCD1 causes depletion of rBMAT of proximal tibia of both sexes, and of cBMAT of distal tibia in male mice only.**

Male and female control and *BMAd-Scd1* KO mice were housed with same-sex littermates and were analyzed at 24-26 weeks of age.

(**A**) Female tibiae were decalcified and stained with osmium tetroxide. Representative bones were rendered in 3D using µCT.

(**B**-**C**) Female tibiae were paraffin-sectioned, stained with hematoxylin and eosin, and photomicrographs obtained for (**B**) proximal tibiae and (**C**) distal tibiae. Scale bar; 200 µm.

(**D**-**E**) MetaMorph software was used to analyze the size of cBMAd in images of distal tibiae.

(**F**) Tibia length was measured at time of sacrifice.

(**G**-**H**) Tibiae were analyzed by µCT for indicated trabecular (Tb.) bone variables. BV/TV: bone volume fraction; BMD: bone mineral density; Conn. Dens: connective density; N: number; Th: thickness; Sp: separation.

(**I**) Tibiae were analyzed by µCT for indicated mid-cortical (Mid Ct.) bone variables. BVF: bone volume fraction.

(**J**-**K**) Tibiae were analyzed by µCT for indicated distal cortical (Dis CT.) bone variables.

(**L**-**M**) Posterior subcutaneous (**L**) and gonadal (**M**) WAT were harvested from male (left) and female (right) mice, paraffin-sectioned, and stained with hematoxylin and eosin. MetaMorph software was used to analyze the size of adipocytes. Data are expressed as mean ± SD.

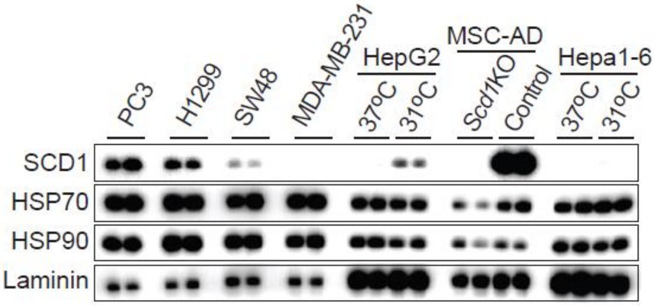


Figure S6

**Supplemental Figure 6. Variable expression of SCD1 explains effects of SCD1 inhibitors on mechanisms of cell death?**

SCD1 expression across the indicated cell lines is demonstrated with immunoblot analysis of 10 µg lysate. All cells, with the exception of adipocytes, maintained sub-confluency (<80%) to avoid hypoxic or nutrient-depleted conditions. Scd1KO adipocytes served as a negative control for SCD1. Cells analyzed include the human prostate cancer cell line (PC3), human non-small cell lung carcinoma cell line (H1299), human colorectal cancer cell line (SW48), triple-negative human breast cancer cell line (MDA-MB-231), human hepatocellular carcinoma cell line (HepG2), MSC adipocytes, and mouse hepatoma cells from C57L/J mice (Hepa1-6). One day post-seeding, the HepG2 and Hepa1-6 cells were cultured at either 37°C or 31°C for a period of 4 days. HSP70, HSP90 and laminin are included as loading controls. Data are representative of 2 independent experiments.

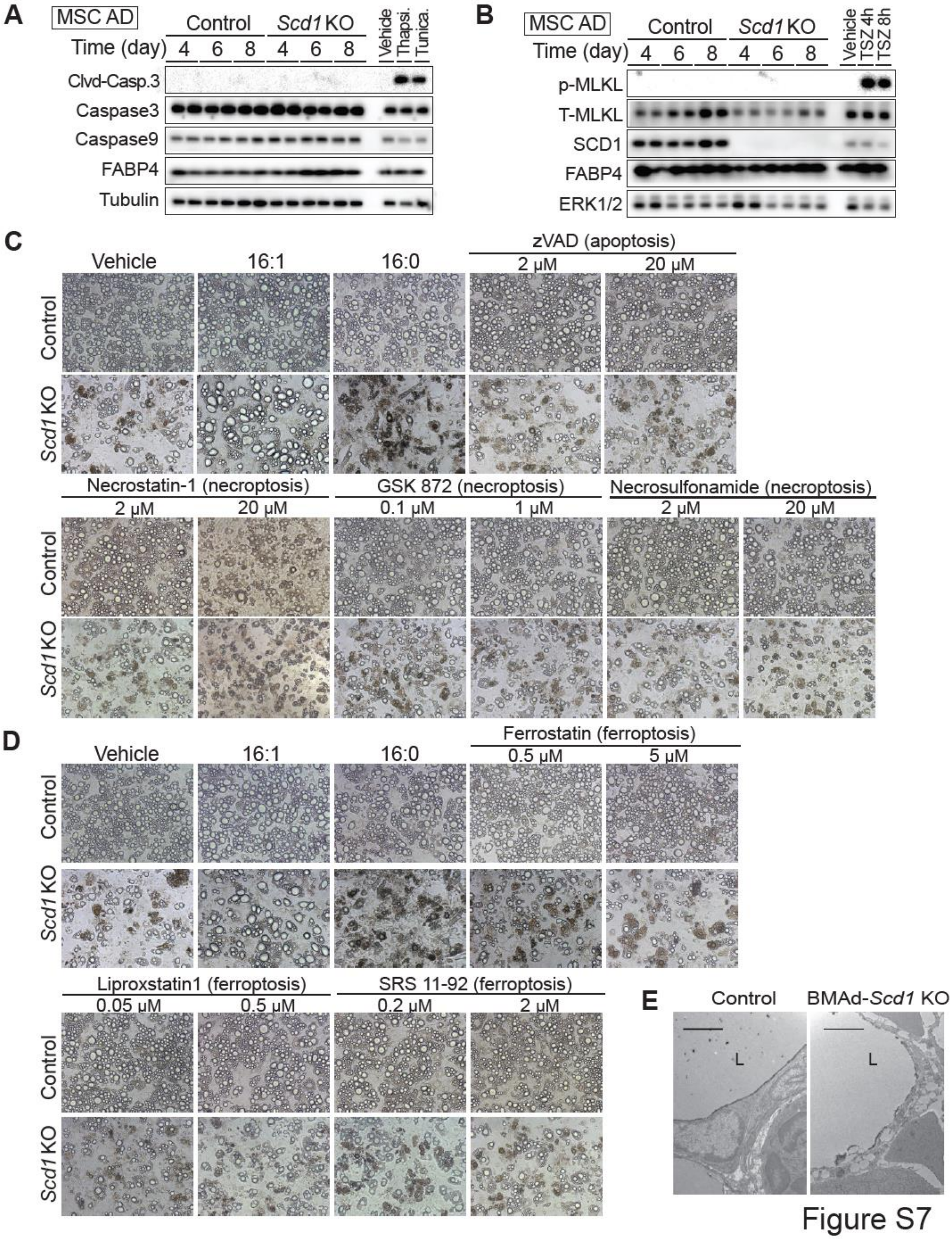

A
MSC AD
Control
Scd1 KO
Time (day)
4
6
8
Vehicle
Thapsi.
Tunica.
Clvd-Casp.3
Caspase3
Caspase9
FABP4
Tubulin
B
MSC AD
Control
Scd1 KO
Time (day)
Vehicle
TSZ 4h
TSZ 8h
p-MLKL
T-MLKL
SCD1
FABP4
ERK1/2
C
Vehicle
16:1
16:0
zVAD (apoptosis)
2 μM
20 μM
Control
Scd1 KO
Necrostatin-1 (necroptosis)
GSK 872 (necroptosis)
0.1 μM
1 μM
Necrosulfonamide (necroptosis)
D
Ferrostatin (ferroptosis)
0.5 μM
5 μM
Liproxstatin1 (ferroptosis)
0.05 μM
SRS 11-92 (ferroptosis)
0.2 μM
2 μM
E
BMAd-Scd1 KO
L


Figure S7

**Supplemental Figure 7 *Scd1*KO adipocytes did not undergo apoptosis nor necroptosis.** (**A** and **B**) Control and *Scd1*KO adipocytes at the indicated days of differentiation were examined by immunoblotting using indicated primary antibodies to detect (**A**) apoptosis or (**B**) necroptosis biomarkers. As a positive control for apoptosis, cultured adipocytes were treated with 1 µM thapsigargin or 1 µM tunicamycin for 24 hrs. As a positive control for necroptosis, cultured adipocytes were treated with mixture of TSZ (20 ng/mL TNF-α, 200 nM SM-164, and 20 µM zVAD-fmk) for 4 or 8 h. Data shown is representative of at least 3 independent experiments.

(**C** and **D**) Control or *Scd1*KO adipocytes were treated with the indicated concentrations of apoptosis inhibitor (zVAD), necroptosis inhibitors (necrostatin-1, GSK872, or necrosulfonamide) (**C**), or ferroptosis inhibitors (ferrostatin, liproxstatin, or SRS11-92) (**D**) from day 4 to 8 of differentiation. Cells were treated with 100 µM palmitoleic acid (C16:1) to rescue viability, or 100 µM palmitic acid (C16:0) acids to potentiate cell death. (**E**) Representative images of distal tibial cBMAT by transmission electron microscopy (TEM). Scale bar; 2 µm. Data shown is representative of 2 independent experiments.
